# Codon-dependent regulation of mRNA translation and stability by ZC3H7A and ZC3H7B RNA-binding proteins

**DOI:** 10.1101/2025.04.21.649502

**Authors:** Patric Harris Snell, Parisa Naeli, Aitor Garzia, Joseph A. Waldron, Susanta Chatterjee, Tom McGirr, Reese Jalal Ladak, Jung-Hyun Choi, Jun Luo, Sami Leino, Naomi Jess, S. Ali Shariati, Ximena Soto Rodriguez, Christos G. Gkogkas, Nahum Sonenberg, Thomas Tuschl, Sarah Maguire, Seyed Mehdi Jafarnejad

**Author notes:** These authors contributed equally to this work (Patric Harris Snell, Parisa Naeli). Corresponding author: Seyed Mehdi Jafarnejad, Patrick G. Johnston Centre for Cancer Research, Queen’s University Belfast, Belfast, BT9 7AE, UK.

## Abstract

Decelerated translation elongation caused by non-optimal codons can reduce mRNA stability through *codon optimality-mediated mRNA degradation*. A key element of this process is the coupling of sensing the mRNA codon usage with the regulation of translation efficiency and stability. We report that two paralog RNA-binding proteins (ZC3H7A and ZC3H7B), which are only found in Chordates, preferentially bind to and reduce the stability and translation of mRNAs enriched in non-optimal codons with A/U at their wobble sites (A/U3 codons). ZC3H7A/B engage with ribosomes that lack elongation factors and induce mRNA degradation or block translation initiation through their interactions with the CCR4-NOT and the GIGYF2/4EHP translation repressor complex, respectively. Depletion of ZC3H7A/B or 4EHP impairs the repression of non-optimal A/U3-rich mRNAs. This study provides insights into a unique mechanism in higher eukaryotes that couples codon usage with the regulation of translation efficiency and mRNA stability.

## Introduction

Rigorous control of mRNA translation and turnover is critical for precise post-transcriptional regulation of gene expression whose dysregulation is linked to a plethora of diseases such as neurological disorders, cancers, and infectious diseases [1, 2]. Furthermore, given recent advances in synthetic mRNA technologies, understanding how cells regulate mRNA translation and turnover has become increasingly important for optimizing the efficacy of mRNA-based therapeutic applications.

Various cis regulatory elements within mRNA can influence its rate of translation and stability. These include codon optimality, defined as the non-uniform decoding rate of certain synonymous codons over others that encode the same amino acid. Synonymous codons are not used equally across the transcriptome [3]. Furthermore, functionally associated and co-regulated mRNAs tend to share similar codon usage patterns [4, 5]. For instance, the synonymous codon usage in human mRNAs that encode proteins associated with the cell cycle is biased towards those with A/U at their wobble sites (A/U3 codons), whereas mRNAs associated with differentiation and pattern-specification prefer G/C3 codons [6, 7].

The presence of non-optimal codons generally leads to slower mRNA translation and enhanced rate of decay, through a loosely defined process named *codon optimality-mediated mRNA degradation* [8, 9]. However, various studies differ on which codons are associated with lower mRNA stability or translation rate [10–12]. In humans, A/U3 codons are enriched in mRNAs with short half-lives, while G/C3 codons are associated with longer mRNA half-life [13], and changes in codon optimality impact the mRNA translation rate and stability [10, 13]. Notably, altering the cellular state can also change the degree by which the codon affects mRNA translation and stability. For instance, during viral infection, which is accompanied by reduced translation, the identity of the codons has a weakened effect on mRNA stability [10]. These variations with cellular state and the fluidity of codon optimality in different biological contexts necessitate the presence of a mechanism(s) that senses the speed at which the ribosomes decode the coding sequence and couples that with the regulation of mRNA translation and decay.

RNA-binding proteins (RBPs) play key roles in translation-dependent mRNA decay [14]. The yeast Dhh1 protein and its mammalian homolog DDX6 have been linked to sensing the translation elongation rate through binding to the ribosome to engender the degradation of non-optimal mRNAs [9, 15–17]. The yeast Not5 subunit of the Ccr4-Not complex and its human homolog CNOT3 bind directly to the vacant ribosomal E-site when translating non-optimal codons and thereby act as a ‘sensor’ of mRNA codon optimality [18, 19]. Importantly, there are significant differences in codon usage and mechanisms of codon-dependent regulation of mRNA translation and decay across different species [14, 20]. While the role of Dhh1 and Not5 in *codon optimality-mediated mRNA degradation* decay in yeast is established, several recent studies in mammalian cells have presented a more complex picture where DDX6 and CNOT3 may be dispensable for either translational repression and/or degradation of non-optimal mRNAs [14, 21–23]. These critical differences underscore the possible presence of other factors in higher eukaryotes that sense non-optimal codons and trigger the *codon optimality-mediated mRNA degradation* process.

Zinc Finger CCCH-Type Containing 7A (ZC3H7A) and its paralog protein ZC3H7B (also known as RoXaN), henceforth collectively referred to as ZC3H7A/B, have emerged in Chordates and associate with RNAs [24] and mRNAs [25, 26]. ZC3H7A is localized to cytoplasmic granules and proximity interaction (BioID) analysis identified proteins related to the regulation of mRNA translation and processing as well as cytoplasmic stress granules among its interactors [27]. Previous studies suggested a role for ZC3H7A/B in the biogenesis of a subset of microRNAs [28] and a PAR-CLIP analysis of RNAs that interact with ZC3H7B revealed its binding to the CDS as well as the 3’ UTR of mRNAs [25]. However, the mechanism(s) by which these paralog proteins regulate microRNA biogenesis and their impacts on the stability or translation of their target mRNAs remain ill-defined. Furthermore, the similarity or differences in mechanism(s) of regulation of target mRNAs between ZC3H7A and ZC3H7B are unknown.

Here, we provide evidence that in contrast to the suggestions in a previous report [28], ZC3H7A/B proteins do not show a tendency for interaction with primary microRNA transcripts (pri-miRNAs). Instead ZC3H7A/B proteins bind to the cytoplasmic mRNAs and regulate their translation and stability in a codon content-dependent manner. Accordingly, ZC3H7A/B proteins preferentially bind to and destabilise the A/U3-rich non-optimal mRNAs. We further demonstrate that ZC3H7A/B interact with CNOT3 and DDX6 as well as translationally inactive (paused) ribosomes that are devoid of translation elongation factors. Finally, we show that ZC3H7A/B proteins also interact with the GIGYF2/4EHP translational repressor complex, which could lead to the translational repression of non-optimal mRNAs.

## Results

### ZC3H7A/B preferentially bind to the CDS and 3’ UTRs of mRNAs

Human ZC3H7A and ZC3H7B encode a 971 and 977 amino acid (aa) protein, respectively. Each protein is predicted to contain an N-terminal tetratricopeptide repeat (TPR) domain, followed by an intrinsically disordered region (IDR), and four (ZC3H7A) or five (ZC3H7B) zinc finger (ZNF) motifs at their C-termini (**Fig. 1A & Supp. Fig. 1A & B**). The two paralog proteins share 47.6% identity in their amino acid sequences. This similarity is higher within the TPR and ZNF domains, whereas the IDRs diverge substantially (**Supp. Fig. 1C**). Notably, these genes have only emerged in Chordates (**Supp. Fig. 2A & B**).

**Figure 1.**
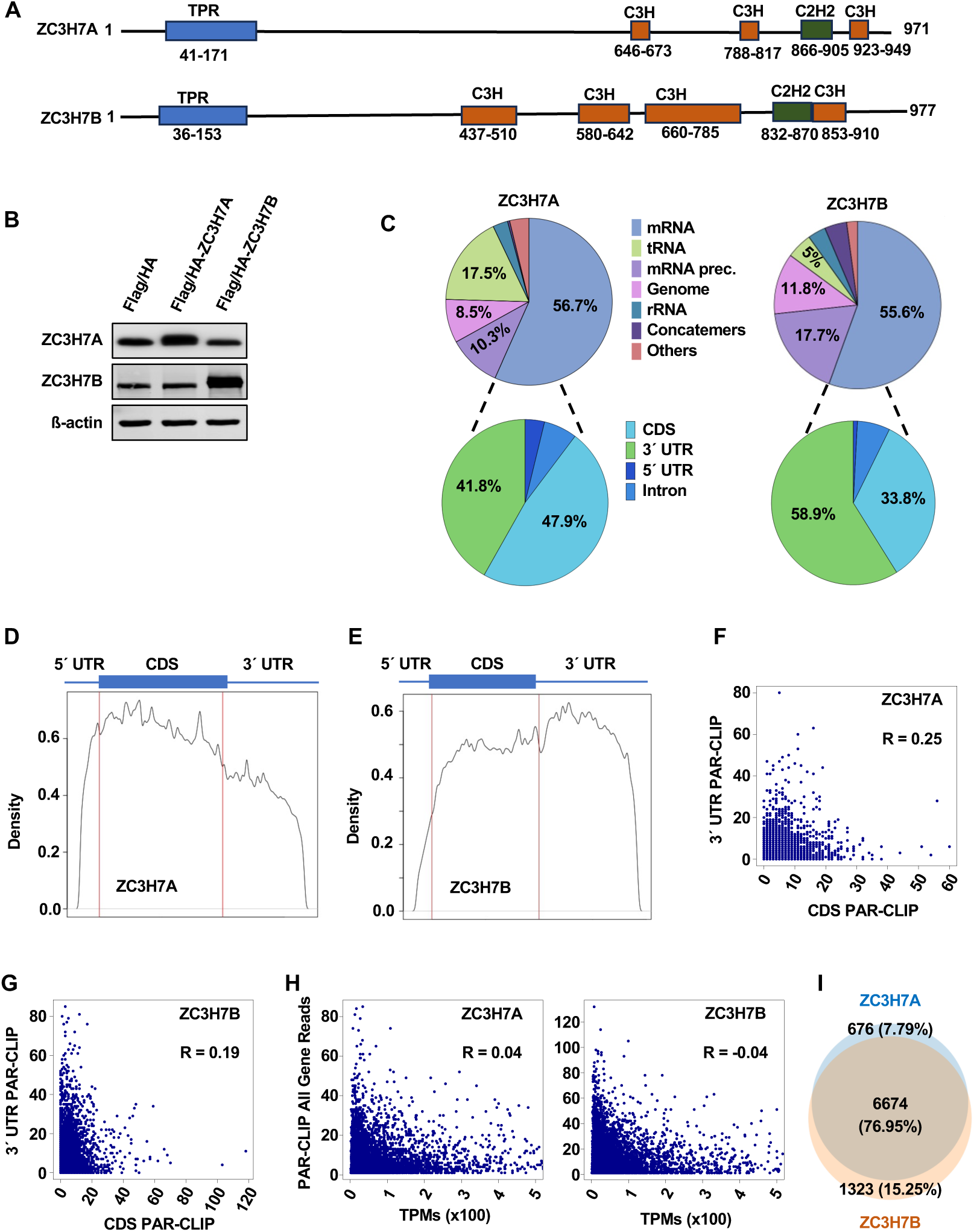
ZC3H7A/B proteins preferentially bind mRNAs within the CDS and 3’ UTR. (**A**) Predicted motifs in human ZC3H7A and ZC3H7B proteins analysed via Interpro (http://www.ebi.ac.uk/interpro/). Numbers indicate the position of amino acids. Domain sizes are not depicted at scale. (**B**) Western blot analysis of the lysates derived from Flag/HA control, Flag/HA-ZC3H7A, and Flag/HA-ZC3H7B in HEK293 cells, 24 h after treatment with doxycycline (1 μg/mL), with the indicated antibodies. (**C**) *Top*: relative composition of ZC3H7A and ZC3H7B PAR-CLIP reads mapping to each RNA category with up to 2 mismatches. *Bottom*: the distribution of binding sites (regions with >50 overlapping reads) across different mRNA regions. (**D & E**) Metagene analysis of the distribution of ZC3H7A (D) and ZC3H7B (E) PAR-CLIP sequence reads across its target mRNAs. (**F & G**) Scatter plot analysis of correlation between the PAR-CLIP reads mapped to CDS vs 3’ UTR of the same mRNAs for ZC3H7A (F) and ZC3H7B (G). R =Pearson correlation coefficient. (**H**) Correlation between the mRNA abundance (RNA-Seq) and the PAR-CLIP reads for the same mRNAs in ZC3H7A-OE and ZC3H7B-OE cells. R =Pearson correlation coefficient. (**I**) Overlap between ZC3H7A and ZC3H7B crosslinked target mRNAs, defined as mRNAs with at least one binding site.

A previous published study concluded that ZC3H7A/B bind the hairpin structures in pri-miRNAs and promote their maturation [28]. High-throughput studies of the mRNA-bound proteome identified both ZC3H7A and ZC3H7B as RBPs that bind to poly-adenylated mRNAs [25, 26] and a PAR-CLIP analysis revealed that ZC3H7B binds to the 3’ UTR and the CDS of select mRNAs [25]. However, the target RNAs of ZC3H7A and the similarity/differences between target RNAs of ZC3H7A and ZC3H7B are unknown. Furthermore, the consequences of the direct binding of ZC3H7A and ZC3H7B to mRNAs is not understood. Therefore, we first sought to identify the transcriptome-wide target RNAs of ZC3H7A and ZC3H7B by performing parallel 4-thiouridine (4SU)-PAR-CLIP assays [29] in HEK293 cells that express a doxycycline-inducible Flag/HA-ZC3H7A (ZC3H7A-OE) or - ZC3H7B (ZC3H7B-OE; **Fig. 1B**). While we did not observe any substantial cross-linking to primary, precursor, or mature microRNAs (**Supp. Table 1**), both ZC3H7A and ZC3H7B mostly cross-linked to mRNAs (56.7% and 55.6% of PAR-CLIP reads, respectively) and to a much lesser extent tRNAs (17.5% and 5%, respectively) (**Fig. 1C & Supp. Table 1**).

While ZC3H7A demonstrated a slight preference for binding to the CDS (47.9% of binding sites; defined as clusters of >50 overlapping reads) compared with 3’ UTR (41.8% of binding sites; **Fig. 1C & D**), ZC3H7B showed a preference for 3’ UTRs (58.9%) compared with the CDS (33.8% of binding sites; **Fig. 1C & E**), which is consistent with previous observations [25]. We did not observe a strong correlation between the binding to the CDS and binding to the 3’ UTR of the same target mRNAs (R =0.25 and 0.19 for ZC3H7A and ZC3H7B, respectively; **Fig. 1F & G**), suggesting that the binding of these proteins to the CDS or 3’ UTR are likely independent events. The cross-linked mRNA read abundance did not resemble the overall mRNA abundance as determined by poly(A) RNA-Seq (R =0.04 and −0.04 for ZC3H7A and ZC3H7B, respectively; **Fig. 1H**), indicating that their binding to mRNAs is likely transcript-specific. We observed a marked overlap between target mRNAs with at least one ZC3H7A or ZC3H7B binding site; while 83% of ZC3H7B target mRNAs also crosslinked to ZC3H7A, 91% of ZC3H7A target mRNAs were also crosslinked to ZC3H7B (**Fig. 1I**). Altogether, these data demonstrate the preferential binding of ZC3H7A/B proteins to the mRNAs across the CDS and 3’ UTR and a significant similarity in their binding patterns and target mRNAs in HEK293 cells.

### ZC3H7A/B proteins preferentially bind to A/U3-rich mRNAs

*De novo* motif analysis revealed the enrichment for an AGAA-rich sequence motif in ZC3H7A and ZC3H7B PAR-CLIP peaks (**Supp. Fig. 3A**). We next investigated the sequence features that were enriched in the target mRNAs. ZC3H7A or ZC3H7B target mRNAs were divided into 5 categories based on the number of binding sites: A =1-2, B =3-4, C =5-9, D =10-24, and E ≥25 (**Supp. Fig. 3B**). The number of binding sites for each protein on the target mRNAs was strongly correlated with the length of the 3’ UTR and only modestly with the length of CDS, but not the 5’ UTR length (**Supp. Fig. 3C-J**). The number of binding sites on the CDS or 3′ UTR was positively correlated with the length of the respective region (**Supp. Fig. 3K-N**) but not with the length of the other region (**Supp. Fig. 3O-R**). Notably, for both proteins we observed a strong negative correlation between the number of binding sites and the G/C content (G/C%) across the CDS and 3’ UTR, but only a weak negative correlation with the 5’ UTR G/C% (**Fig. 2A-H**). We next assessed if these proteins have a bias for binding to mRNAs enriched for non-optimal A/U3 codons or the optimal G/C3 codons. We observed a marked preference in the binding of ZC3H7A and ZC3H7B to A/U3-rich mRNAs, compared with G/C3-rich mRNAs (**Fig. 2I & J**). This correlation was even more pronounced for the CDS binding sites (**Supp.** Fig. 4A **& B**), compared with those mapped to 3’ UTR (**Supp.** Fig. 4C **& D**). To further corroborate these findings, we performed a similar analysis of a previously published ZC3H7B PAR-CLIP data [25], which revealed a remarkably similar preference in binding to A/U3-rich mRNAs (**Fig. 2K**).

**Figure 2.**
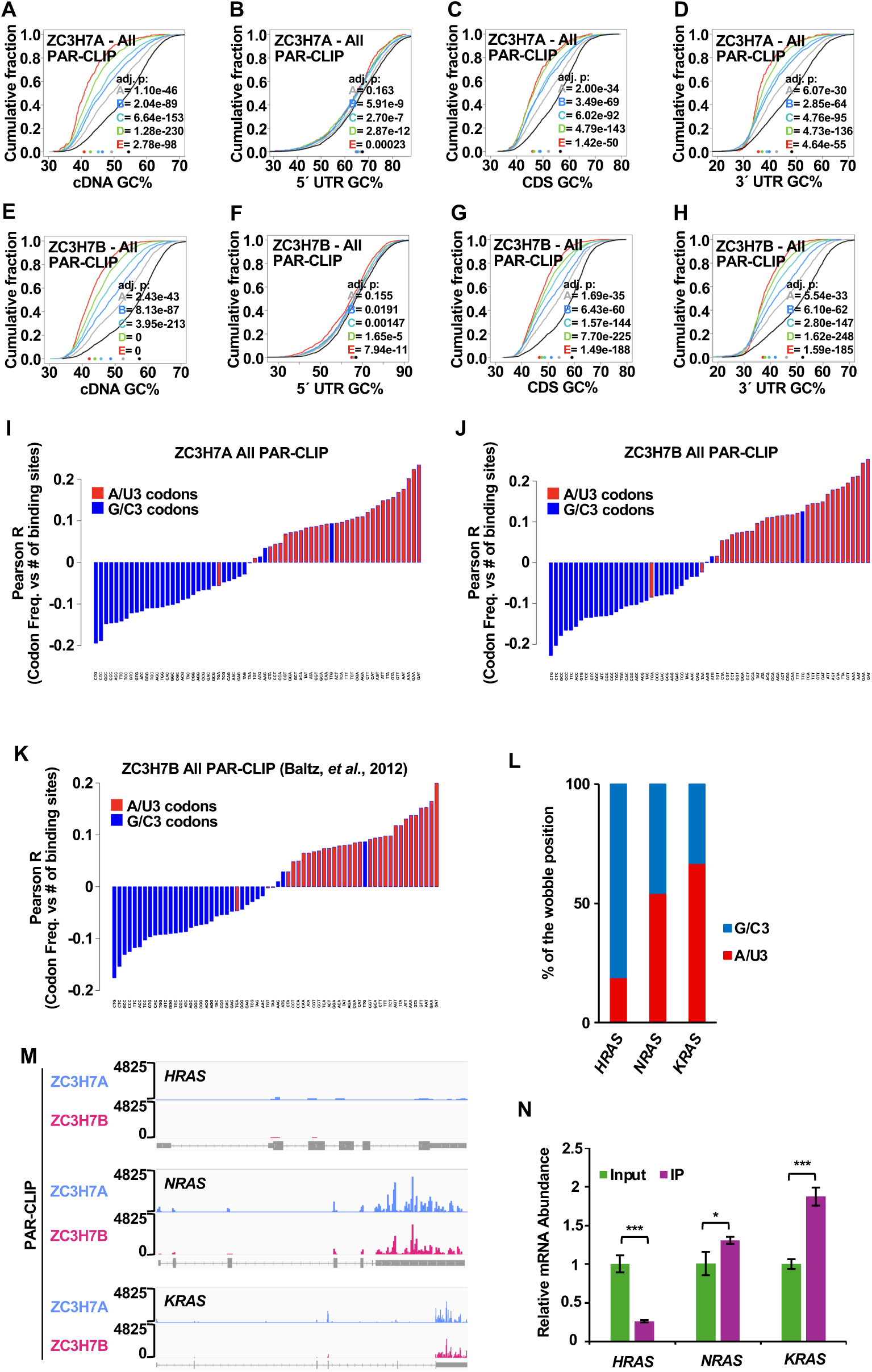
Preferential binding of ZC3H7A/B proteins to non-optimal A/U3 rich mRNAs. (**A-H**) Cumulative distribution analysis of the correlation between the G/C content (G/C%) of the indicated mRNA regions and the number of ZC3H7A or ZC3H7B binding sites. Target mRNAs were divided into 5 categories based on the number of binding sites; A =1-2, B =3-4, C =5-9, D =10-24, and E ≥25. Numbers represent the adjusted p-values (Dunn test) of comparisons between the indicated group and the control (0 binding site). (**I & J**) Waterfall plots showing Pearson correlations between codon usage frequencies and number of ZC3H7A (I) and ZC3H7B (J) PAR-CLIP binding sites. (**K**) Waterfall plot showing Pearson correlations between codon usage frequencies and number of ZC3H7B binding sites in HEK293 cells obtained from Baltz, *et al.* [25]. (**L**) Percentage of A/U3 and G/C3 codons in human *HRAS*, *NRAS* and *KRAS* CDS. (**M**) IGV tracks of ZC3H7A and ZC3H7B PAR-CLIP reads for the *HRAS*, *NRAS* and *KRAS* mRNAs. Gene sizes are not depicted at scale. (**N**) RNA-IP and RT-qPCR analysis of the association between Flag/HA-ZC3H7B and *HRAS*, *NRAS,* and *KRAS* mRNAs in ZC3H7B-OE cells. Data are presented as mean ± SD (n =3). \**P* <0.05, \*\*\**P* <0.001; unpaired t-test.

We validated our findings by comparing the binding of ZC3H7A/B to mRNAs encoded by members of the *RAS* family (*HRAS*, *NRAS*, and *KRAS*). Despite the high level (∼85%) of homology in their amino acid sequences [30], their mRNAs markedly vary in their codon usage with *HRAS* encoding the most optimal mRNA (81% G/C3), followed by *NRAS* (46% G/C3), and *KRAS* (33% G/C3; **Fig. 2L & Supp. Fig. 4E**). Our RNA-Seq analysis (described below) showed that these genes are expressed at relatively similar levels in HEK293 cells (**Supp. Fig. 4F**). However, while we did not detect any PAR-CLIP binding site for either ZC3H7A or ZC3H7B on the *HRAS* mRNA, ZC3H7A and ZC3H7B binding sites mapping to both the CDS and 3’ UTRs of non-optimal *NRAS* and *KRAS* mRNAs were highly abundant (**Fig. 2M**). We further confirmed these results by RNA-Immunoprecipitation (RNA-IP) assay, which revealed a significant enrichment for the non-optimal *KRAS*, along with a modest enrichment for *NRAS*, but no enrichment for the optimal *HRAS* mRNA in ZC3H7B-bound RNAs (**Fig. 2N & Supp. Fig. 4G**). Together, these data demonstrate that the binding of ZC3H7A/B to the mRNAs is strongly influenced by the abundance of A/U3 codons.

### ZC3H7A/B binding leads to destabilisation of A/U3-rich mRNAs

The strongly biased binding of the ZC3H7A/B proteins to the A/U3-rich non-optimal mRNAs suggests that these proteins may play a role in the *codon optimality-mediated mRNA degradation* process [8, 9]. To identify the effects of ZC3H7A and ZC3H7B binding to the mRNAs, we performed poly(A) RNA-Seq and Ribo-Seq (ribosome profiling) on samples derived from ZC3H7A-OE and ZC3H7B-OE cells, compared with the Flag/HA expressing control (CTRL) HEK293 cells (**Supp. Fig. 5A-C**). We observed substantial changes in mRNA expression and translation upon overexpression of ZC3H7A or ZC3H7B. 3512 and 2968 mRNAs were downregulated while 3134 and 2087 mRNAs were upregulated in the ZC3H7A-OE and ZC3H7B-OE cells respectively, compared with the CTRL cells (FC>1.5; Adj-p <0.05; **Fig.** 3A & B **and Supp. Table 2**). Similarly, the ribosome occupancy [read depth-normalised number of ribosome-protected fragments (RPFs)] of 1340 and 1237 mRNAs were downregulated while 1601 and 1277 mRNAs were upregulated in the ZC3H7A-OE and ZC3H7B-OE cells respectively (FC>1.5; Adj-p <0.05; **Fig. 3C & D and Supp. Table 2**). Importantly, consistent with the high level of similarity in the PAR-CLIP target mRNAs of ZC3H7A and ZC3H7B (**Fig. 1I**), we observed a high degree of similarity in the impact of ZC3H7A-OE and ZC3H7B-OE on mRNA abundance (Spearman ϱ =0.78; **Fig. 3E**) and ribosome occupancy (ϱ =0.72; **Fig. 3F**). This reflects a high degree of redundancy in their target mRNAs, functions, and likely biological impacts. In addition, we observed a positive correlation between the changes in mRNA abundance and ribosome occupancy in both ZC3H7A-OE (ϱ =0.46; **Supp. Fig. 5D**) and ZC3H7B-OE cells (ϱ =0.42; **Supp. Fig. 5E**). This suggests that ZC3H7A/B impact the translation and stability of the target mRNAs mostly in similar directions.

**Figure 3.**
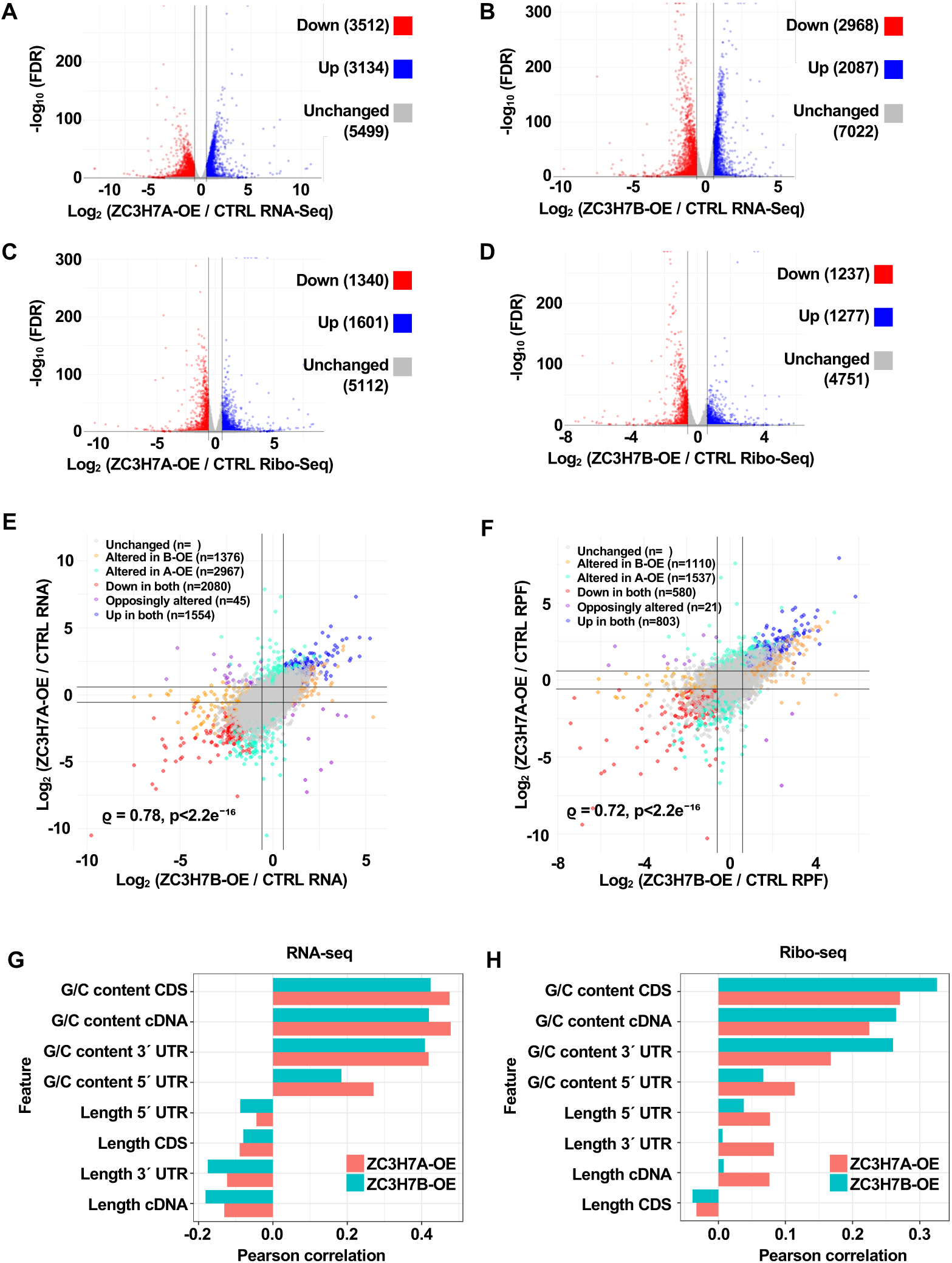
Dysregulated mRNA expression and translation upon ZC3H7A/B overexpression. (**A & B**) Volcano plots showing differentially expressed (FC >1.5; FDR <0.05) mRNAs in ZC3H7A-OE (A) or ZC3H7B-OE (B) compared with the Flag/HA control cells. (**C & D**) Volcano plots showing mRNAs with differential (FC >1.5; FDR<0.05) ribosome occupancy in ZC3H7A-OE (C) or ZC3H7B-OE (D) compared with the Flag/HA control cells. (**E & F**) Scatter plot showing the comparative analysis of differential (FC >1.5; FDR <0.05) mRNA abundance (E) or ribosome occupancy (F) in ZC3H7A-OE and ZC3H7B-OE cells compared with the Flag/HA control cells. ϱ =Spearman correlation coefficient. (**G & H**) Comparison of Pearson correlations between the indicated mRNA features and the changes in mRNA expression (G) or ribosome occupancy (H) in ZC3H7A-OE or ZC3H7B-OE cells. Only mRNAs with FDR <0.05 were included and the most abundant transcript per gene (based on total RNA-seq data) was selected.

Upon analysis of the mRNA sequence features, we observed a weak or no correlation between the length of the cDNA, CDS, and UTRs and changes in the mRNA abundance (**Fig. 3G & Supp. Fig. 6A-H**) or ribosome occupancy (**Fig. 3H & Supp. Fig. 6I-P**) in the ZC3H7A-OE and ZC3H7B-OE cells compared with CTRL cells. In contrast, the changes in mRNA expression and ribosome occupancy were strongly and positively correlated with the mRNA G/C% (**Fig. 3G & H and Supp. Fig. 7A-P**). Conspicuously, these correlations were markedly stronger when considering the G/C% of the CDS and 3’ UTR, compared with the 5’ UTR. Furthermore, mRNAs that were upregulated in ZC3H7A-OE and ZC3H7B-OE (**Fig. 4A & B**) or mRNAs with increased ribosome occupancy (**Supp. Fig. 8A & B**) were highly enriched in G/C3 codons, whereas the downregulated mRNAs were highly enriched in A/U3 codons. Notably, TTG, one of the 6 Leucine codons defied this pattern and was the only G/C3 codon enriched among downregulated mRNAs (**Fig. 4A & B** and **Supp.** Fig 8A **& B**). TTG was also the only G/C3 codon enriched in ZC3H7A/B PAR-CLIP target mRNAs (**Fig. 2I-K**). Interestingly, previous studies revealed that TTG deviates from and shows an inverse relationship with G/C3 content [31–33].

**Figure 4.**
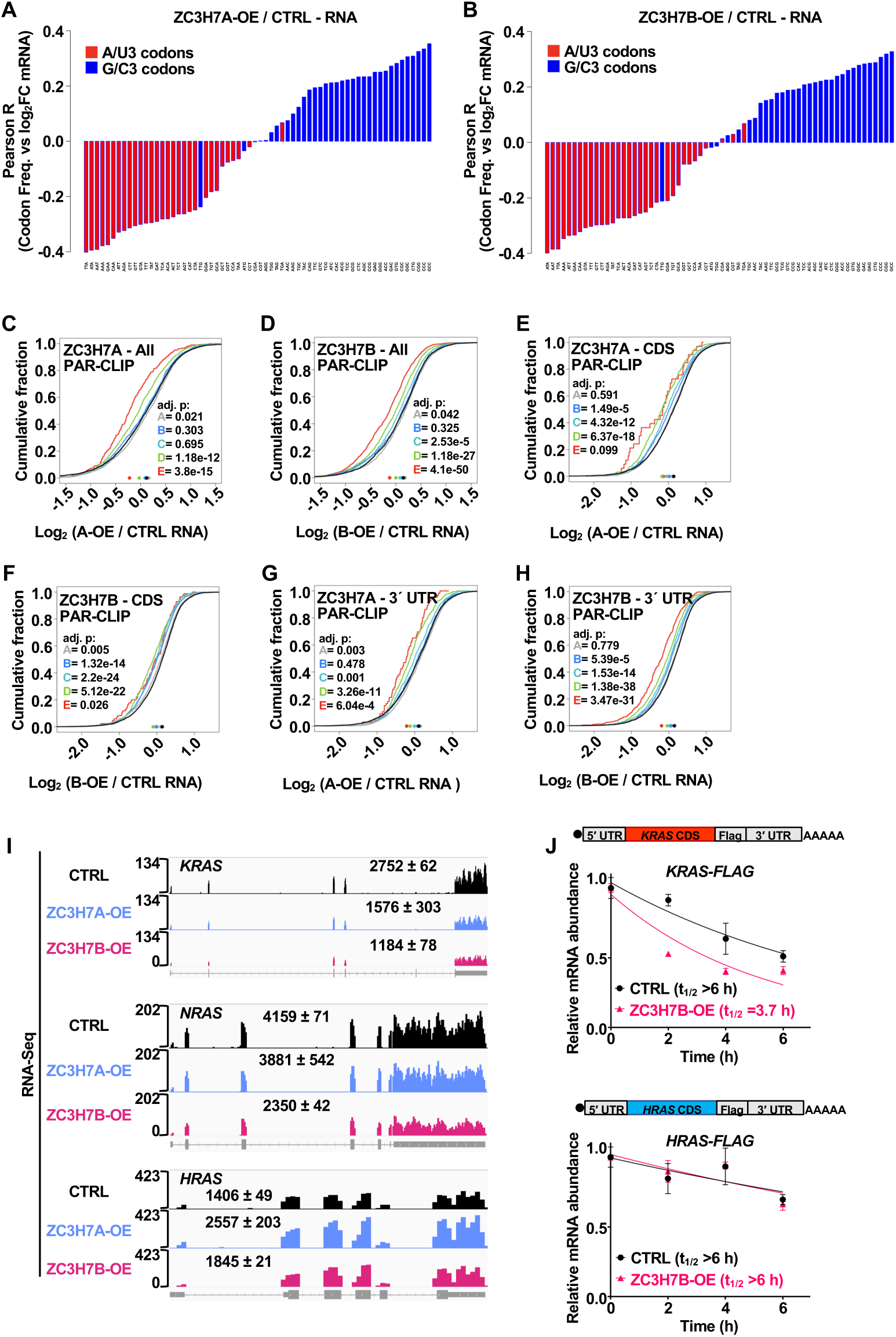
Divergent impacts of ZC3H7A/B on expression of optimal vs non-optimal mRNAs. (**A & B**) Waterfall plots showing Pearson correlations between codon usage frequencies and changes in mRNA expression in ZC3H7A-OE (A) and ZC3H7B-OE (B) compared with Flag/HA control cells. (**C-H**) Cumulative distribution analysis of the number of ZC3H7A or ZC3H7B binding sites across the indicated mRNA regions of differentially expressed mRNAs (FC >1.5; FDR <0.05) in ZC3H7A-OE or ZC3H7B-OE compared to Flag/HA control cells. (**I**) IGV tracks visualisation of *HRAS*, *NRAS* and *KRAS* mRNAs in the Flag/HA control, ZC3H7A-OE, and ZC3H7B-OE cells. The normalized RNA count for each gene is presented as mean ± SD. (**J**) RT-qPCR analysis of the half-life (t_1/2_) of *in vitro* transcribed *KRAS-FLAG* (Top) and *HRAS-FLAG* (Bottom) mRNAs in Flag/HA control and ZC3H7B-OE cells.

We next investigated the relationship between the ZC3H7A/B proteins’ binding to mRNAs (PAR-CLIP) and the changes in the mRNA expression. We noted a negative correlation between the number of binding sites for either ZC3H7A (**Fig. 4C**) or ZC3H7B (**Fig. 4D**) and the changes in the mRNA expression in the corresponding cells. Similar negative correlations were observed when comparing the changes in mRNA expression and number of binding sites in CDS (**Fig. 4E & F**) and 3’ UTR (**Fig. 4G & H**). Notably, unlike the codon-optimal *HRAS* mRNA, expression (**Fig. 4I**) and ribosome occupancy (**Supp. Fig. 8C**) of the non-optimal *KRAS*, and to a lesser extent *NRAS* mRNA, were downregulated in ZC3H7A-OE and ZC3H7B-OE cells compared with the control cells. To confirm that the divergent effects on the *HRAS* and *KRAS* mRNAs was due to the differences in their CDS codon usage and not some other features, particularly in their UTRs, we generated synthetic mRNAs with identical 5’ and 3’ UTRs and either the *HRAS* or *KRAS* CDS, which were tagged with an in-frame Flag epitope at their C-terminus. The *in vitro* transcribed mRNAs were transfected into the ZC3H7B-OE or Flag/HA control cells. As expected, ZC3H7B-OE resulted in substantially reduced stability of the non-optimal *KRAS-FLAG* mRNA but not the optimal *HRAS-FLAG* mRNA (**Fig. 4J**). Altogether, these data demonstrate that ZC3H7A/B binding leads to destabilisation of the non-optimal A/U3-rich mRNAs.

### Depletion of the ZC3H7A/B proteins impairs the translation-dependent decay of non-optimal mRNAs

We next studied the impact of depletion of ZC3H7A and ZC3H7B proteins on mRNA translation and stability. We reasoned that, considering the marked redundancies in the target mRNAs of ZC3H7A and ZC3H7B in HEK293 cells, as determined by the PAR-CLIP (**Fig. 1I**), RNA-Seq (**Fig. 3E**), and Ribo-Seq assays (**Fig. 3F**), simultaneous depletion of the two proteins is required for sensitive assessment of their impacts on mRNA translation and stability. However, while we readily achieved CRISPR-Cas9 mediated knockout (KO) of either gene in HEK293 cells (**Supp. Fig. 9A**), we failed to generate a cell line wherein the two genes were simultaneously depleted via CRISPR-Cas9. Thus, we used shRNA to knockdown (KD) ZC3H7B expression in ZC3H7A-KO cells (**Supp. Fig. 9B**), which we dubbed ZC3H7A/B–double depletion (ZC3H7A/B-DD) cells. Of note, while CRISPR-mediated depletion of individual ZC3H7A or ZC3H7B proteins exerted only small effects on cell growth (**Supp. Fig. 9C**), ZC3H7A/B-DD cells grew at markedly lower rate than the Parental cells expressing a non-targeting control shRNA (**Supp. Fig. 9D**).

RNA-Seq and Ribo-Seq analyses of the ZC3H7A/B-DD compared with the Parental cells (**Supp. Fig. 9E-G and Supp. Table. 3**) revealed substantial changes in mRNA expression and ribosome occupancy. The abundance of 598 mRNAs was upregulated and 1043 mRNAs were downregulated (FC >1.5; Adj-p <0.05; **Fig. 5A**), while ribosome occupancy of 863 mRNAs was increased, and 1247 mRNAs were decreased (FC >1.5; Adj-p <0.05; **Fig. 5B**) in ZC3H7A/B-DD cells compared with Parental cells. Comparative analysis with the ZC3H7A/B PAR-CLIP data revealed a statistically significant, albeit modest positive correlation between the number of ZC3H7A or ZC3H7B binding sites and changes in mRNA expression (**Fig. 5C & D**) and ribosome occupancy (**Fig. 5E & F**) in ZC3H7A/B-DD cells. As expected, this pattern is the opposite of the observed negative correlation between the number of binding sites and differentially expressed mRNAs in ZC3H7A-OE and ZC3H7B-OE cells (**Fig. 4C-H**). We next examined the correlation between changes in mRNA expression and ribosome occupancy with ZC3H7A/B binding to CDS vs 3’ UTR. The positive correlations were even more prominent with the number of binding sites in the CDS (**Fig. 5G-J**) compared with 3’ UTR (**Supp. Fig. 9H-K**). Consistent with these observations, analysis of codon usage bias in differentially expressed (**Fig. 5K**) or translated (**Fig. 5L**) mRNAs in ZC3H7A/B-DD cells identified an enrichment for A/U3 codons in upregulated mRNAs and G/C3 codons in down-regulated mRNAs, the inverse to what was observed in ZC3H7A-OE and ZC3H7B-OE cells (**Fig. 4A-B & Supp. Fig. 8A-B**). We next used a reporter assay [34] wherein synonymous codons are used to generate mCherry coding mRNAs with different percentages (100% vs 25%) of G/C3 codons to assess the impact of the depletion of ZC3H7A/B on the reporters’ expression, in relation to their codon usage. While expression of the optimal (100% G/C3) reporter decreased by 24%, expression of the non-optimal (25% G/C3) reporter increased by 37% in ZC3H7A/B-DD cells, compared with the Parental cells (p <0.001; **Fig. 5M**). Collectively, these data suggest that depletion of the ZC3H7A/B proteins impairs the codon-optimality mediated regulation of gene expression.

**Figure 5.**
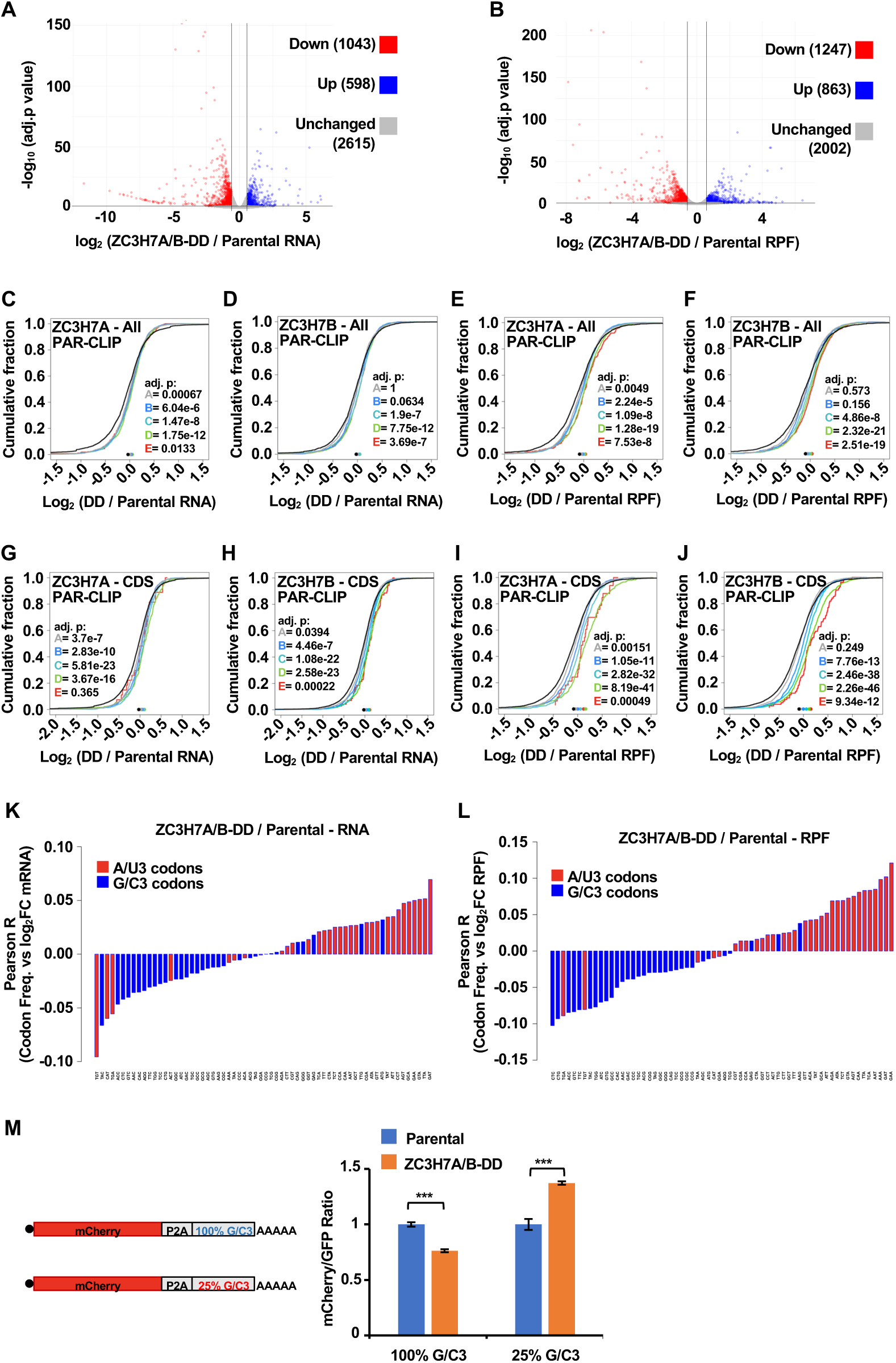
Impairment of translation-dependent decay mechanism in ZC3H7A/B-depleted cells. (**A & B**) Volcano plots showing differential (FC >1.5; FDR <0.05) mRNA expression (A) and ribosome occupancy (B) in ZC3H7-DD compared to the parental HEK293 cells transduced with a non-targeting control shRNA (Parental). (**C & D**) Cumulative distribution analysis of the correlation between the changes in mRNA expression in ZC3H7A/B-DD compared to the Parental cells and the number of ZC3H7A (C) and ZC3H7B (D) PAR-CLIP binding sites across all regions of the mRNAs. (**E & F**) Correlation between the changes in ribosome occupancy in ZC3H7A/B-DD compared to Parental cells and the number of ZC3H7A (E) and ZC3H7B (F) PAR-CLIP binding sites across all regions of the mRNAs. (**G & H**) Correlation between the changes in mRNA expression in ZC3H7A/B-DD compared to Parental cells and the number of ZC3H7A (G) and ZC3H7B (H) binding sites in CDS. (**I & J**) Correlation between the changes in ribosome occupancy in ZC3H7A/B-DD compared to Parental cells and the number of ZC3H7A (I) and ZC3H7B (J) binding sites in CDS. (**K & L**) Waterfall plots showing Pearson correlations between codon usage frequencies and changes in mRNA expression (K) and ribosome occupancy (L) in ZC3H7A/B-DD cells compared with Parental cells. (**M**) Quantification of flow cytometry measurement of the fluorescence intensity of 100% and 25% G/C3 mCherry reporter, normalised to the GFP control, 24 h after transfection into ZC3H7A/B-DD and Parental cells. Data are presented as mean ± SD (n =3). \*\*\**P* <0.001; unpaired t-test.

### ZC3H7A/B binding leads to CCR4-NOT-dependent mRNA degradation

We next sought to identify the mechanism(s) by which ZC3H7A/B binding to the 3’ UTR affects target mRNAs by using the LambdaN (λN):BoxB tethering system [35], with a *Renilla* Luciferase (RL) reporter that contains five BoxB hairpins in its 3’ UTR (RL-5BoxB; **Fig. 6A**). The reporter was co-transfected into HEK293 cells along with a plasmid encoding the ZC3H7A or ZC3H7B protein fused to λN peptide (**Supp. Fig. 10A**), which binds to the BoxB hairpin. A Firefly luciferase (FL) reporter without BoxB hairpin was also co-transfected as a control. Direct binding of λN-ZC3H7A and λN-ZC3H7B reduced the RL-5BoxB reporter activity by 57% (p <0.001) and 45% (p <0.01) respectively compared to the λN-Empty control (**Fig. 6A**). Measurement of the RL reporter mRNA levels by RT-qPCR demonstrated a 40% and 30% reduction upon binding of λN-ZC3H7A and λN-ZC3H7B, respectively (**Fig. 6B**). Furthermore, measurement of the RL reporter mRNA stability after blocking new transcription by Actinomycin D showed a marked reduction in mRNA half-life upon tethering with λN-ZC3H7A (1.4 h) or λN-ZC3H7B (2.6 h), when compared to the λN control (>4 h; **Fig. 6C**). These data support our integrative analysis of the PAR-CLIP and RNA-Seq data, which showed the ZC3H7A/B binding to the 3’ UTR correlates with reduced mRNA levels (**Fig. 4G & H**).

**Figure 6.**
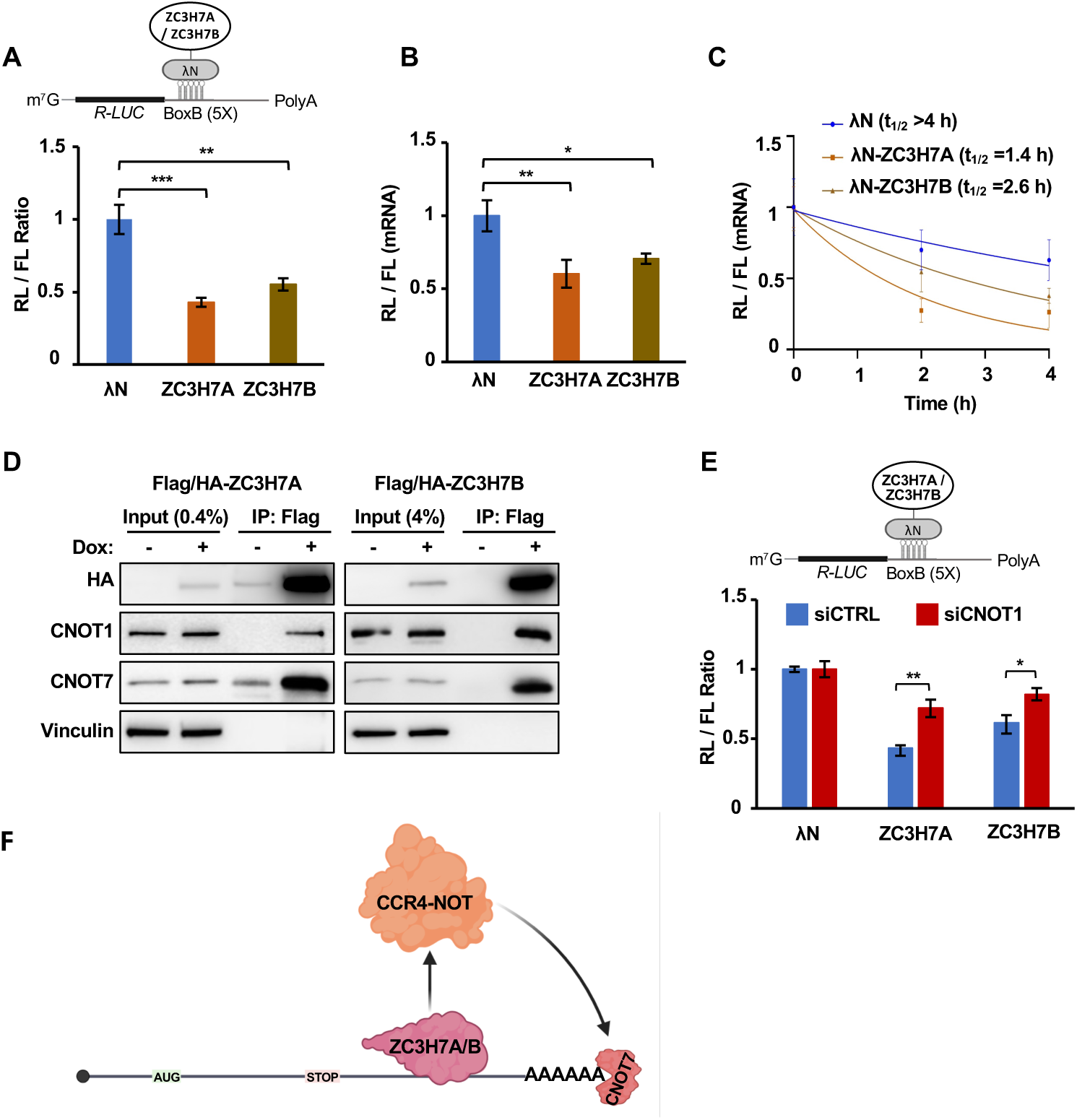
CCR4-NOT-mediated mRNA degradation upon ZC3H7A/B binding to 3’ UTR. (**A**) *Top*: schematic representation of the *RL*-5BoxB reporter. *Bottom*: Analysis of the relative silencing of the *RL*-5BoxB reporter upon tethering with λN-empty, λN-ZC3H7A, or λN-ZC3H7B in HEK293 cells. Data are presented as mean ± SD (n =3). (**B**) RT-qPCR analysis of *RL*-5BoxB mRNA relative to FL mRNA expression in samples derived from (A). Data are presented as mean ± SD (n =3). (**C**) RT-qPCR analysis of *RL*-5BoxB mRNA half-life (t_1/2_) after transfection with *RL*-5BoxB reporter and either λN-empty, λN-ZC3H7A, or λN-ZC3H7B, along with the FL plasmid (control) followed by transcription inhibition using Actinomycin D. Data are presented as mean ± SD (n =3). (**D**) Co-IP for detection of interaction between Flag/HA-ZC3H7A or Flag/HA-ZC3H7B and indicated proteins in HEK293 cells. IPs were performed in presence of RNase A. (**E**) Analysis of the relative silencing of the *RL*-5BoxB reporter upon tethering with λN-empty, λN-ZC3H7A, or λN-ZC3H7B in CNOT1-depleted HEK293 cells. Cells were transfected with siCTRL or siCNOT1 and 48 h later were co-transfected with *RL*-5BoxB reporter and either λN-empty, λN-ZC3H7A, or λN-ZC3H7B, along with the *FL* plasmid (control). Cells were lysed after 24 h and luciferase activity was measured. Data are presented as mean ± SD (n =3). \**P* <0.05, \*\**P* <0.01, \*\*\**P* <0.001; two-tailed unpaired Student’s *t*-test. (**F**) Graphic illustration of the proposed mechanism by which ZC3H7A/B mediate mRNA decay upon binding to the mRNA 3’ UTR through recruitment of the CCR4-NOT complex, including the deadenylase subunit CNOT7. Image was generated in BioRender.

Published protein proximity analysis of ZC3H7A via BioID [27, 36] showed interactions with components of the CCR4-NOT complex, which is frequently recruited by RBPs to cause mRNA degradation [37]. Conversely, BioID analyses of the CCR4-NOT subunits identified ZC3H7A and ZC3H7B among interactors of its core subunit CNOT1 and the deadenylase subunits CNOT7 and CNOT6L [36]. Our co-immunoprecipitation (co-IP) assay confirmed the interactions between ZC3H7A/B and CNOT1 as well as CNOT7 (**Fig. 6D**). Furthermore, siRNA-mediated reduction of CNOT1 expression (**Supp. Fig. 10B**), markedly diminished the repression of the RL-5BoxB reporter by either λN-ZC3H7A (49% reduction) or λN-ZC3H7B (46% reduction; **Fig. 6E**). These data demonstrate that the direct binding of ZC3H7A/B promotes mRNA degradation in a CCR4-NOT-dependent manner (**Fig. 6F**).

### ZC3H7A/B interact with paused ribosomes

Previous reports have revealed the key roles of the CNOT3 [21, 38] and DDX6 [15, 16] proteins in *codon optimality-mediated mRNA degradation*. Both CNOT3 and DDX6 have been identified as ZC3H7A proximity interactors [27, 36]. We confirmed these interactions by co-IP (**Fig. 7A**). DDX6 and CNOT3 physically bind the ribosomes, likely to sense the translation speed [15, 16, 18]. When a ribosome encounters a non-optimal codon, the E-site tRNA may be released before the A-site codon is decoded, enabling CNOT3 to enter the vacant E-site [18, 21, 38]. We found that besides CNOT3, immunoprecipitation with ZC3H7A/B also results in co-precipitation of proteins from both large (RPL3) and small (RACK1) ribosomal subunits, but not the translation elongation factors eEF1A or eEF2 (**Fig. 7B**). This suggests that ZC3H7A/B engage with translationally inactive (paused) ribosomes that lack elongation factors, a situation which is anticipated upon decoding of non-optimal codons by ribosomes. Congruently, western blot analysis of the distribution of ZC3H7A/B proteins in sucrose gradients showed that the majority of either protein fractionates in light sucrose gradient fractions, which correspond to poorly translated mRNAs (**Supp. Fig. 11A & B**). Notably, siRNA-mediated depletion of either CNOT3 or DDX6, did not affect the co-precipitation of ribosomal proteins with ZC3H7B (**Fig. 7C**), suggesting that these interactions with translationally inactive ribosomes are independent of CNOT3 and DDX6.

**Figure 7.**
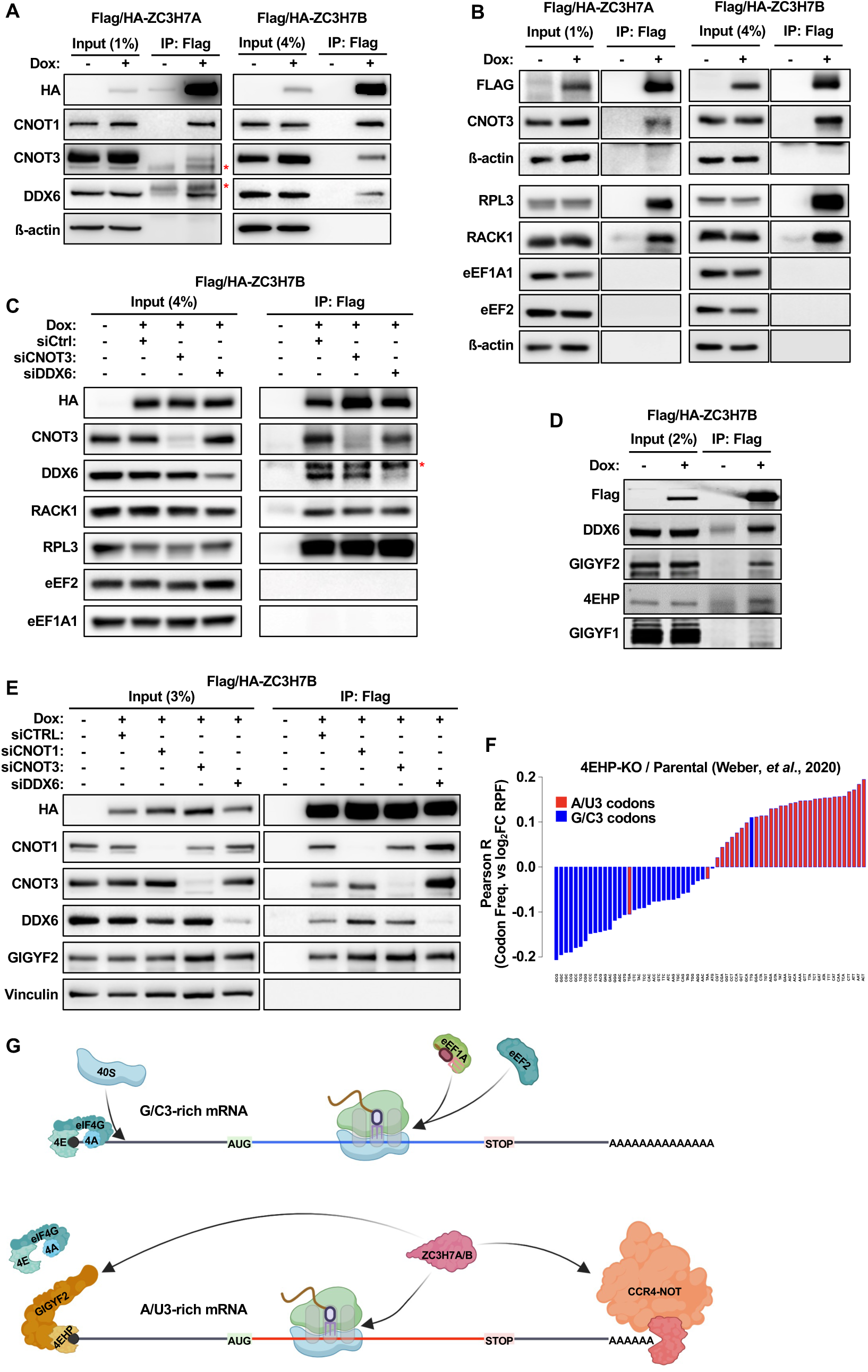
ZC3H7A/B interact with the translationally inactive ribosomes and the GIGYF2/4EHP translation initiation repressors. (**A & B**) Co-IP for detection of interaction between Flag/HA-ZC3H7A or Flag/HA-ZC3H7B and indicated proteins in HEK293 cells. 24 h after doxycycline-induced expression of Flag/HA-ZC3H7A or Flag/HA-ZC3H7B in ZC3H7A-KO or ZC3H7B-KO cells respectively, proteins were immunoprecipitated using the anti-Flag antibody and western blotting was performed with the specified antibodies. *Marks the non-specific bands detected by the antibody. All IP assays were performed in presence of RNase A. (**C**) Co-IP for detection of interaction between Flag/HA-ZC3H7B and indicated proteins in CNOT3-knockdown or DDX6-knockdown ZC3H7B-KO cells. (**D**) Co-IP for detection of interaction between Flag/HA-ZC3H7B and indicated proteins 24 h after doxycycline-induced expression of Flag/HA-ZC3H7B in ZC3H7B-KO cells. (**E**) Co-IP for detection of interaction between Flag/HA-ZC3H7B and indicated proteins in CNOT1-knockdown, CNOT3-knockdown, and DDX6-knockdown ZC3H7B-KO cells. (**F**) Waterfall plots showing Pearson correlations between codon usage frequencies and changes in ribosome occupancy in 4EHP-knockout HEK293 cells compared with Parental cells (obtained from Weber *et al.* [42]). (**G**) The proposed mechanism of translation-dependent regulation of non-optimal A/U3-rich mRNAs by ZC3H7A/B proteins. ZC3H7A or ZC3H7B interact with non-optimal A/U3-rich mRNAs and the ribosomes that lack eEF2 and eEF1A; elongation factors that are typically found on actively translating ribosomes. Binding of ZC3H7A/B to the non-optimal mRNAs may result in repression of new rounds of translation through recruitment of the GIGYF2/4EHP repressor complex or induction of mRNA degradation via CCR4-NOT. Image was generated in BioRender.

A recent study demonstrated that reduced binding of the eIF4F complex, including the cap-binding protein eIF4E, is required for repression of translation initiation on non-optimal mRNAs via a mechanism that is independent of CNOT3 and DDX6 [22]. The cap-binding protein 4EHP (eIF4E Homologous Protein; also known as eIF4E2) represses mRNA translation by displacing eIF4E from the cap [39, 40]. 4EHP is recruited to the mRNAs via RBPs such as Tristetraprolin (TTP) [41], microRNA-Induced Silencing Complex (miRISC) [39], and the Ribosome Quality Control (RQC) factors ZNF598 and EDF1 [42–44]. The Grb10-interacting GYF protein 2 (GIGYF2) acts as a cofactor by bridging 4EHP to the RBPs [39, 41–44]. Notably, both 4EHP and GIGYF2 have been identified as ZC3H7A proximity interactors [27, 36]. Co-IP confirmed that GIGYF2 and 4EHP interact with ZC3H7A/B (**Fig. 7D & Supp. Fig. 11C**). RNAi-mediated depletion of CNOT1, CNOT3, or DDX6 did not abrogate this interaction (**Fig. 7E**). This indicates that the interactions of the ZC3H7A/B proteins and the GIGYF2/4EHP translation initiation repressor complex are independent of the CCR4-NOT complex or DDX6. To establish a role for the GIGYF2/4EHP complex in codon-dependent translational regulation we investigated a published dataset [42] for the potential bias for non-optimal A/U3 or the optimal G/C3 codons among the differentially translated mRNAs in 4EHP-knockout HEK293 cells. Consistent with our observation in ZC3H7A/B-DD cells (**Fig. 5L**), the translationally upregulated mRNAs in 4EHP-KO cells were enriched with A/U3 codons, whereas the downregulated mRNAs were enriched with G/C3 codons (**Fig. 7F**).

Altogether, these data indicate that besides CNOT3, DDX6, and ribosomes that lack elongation factors, ZC3H7A/B proteins also interact with the GIGYF2/4EHP, which could in turn displace the eIF4F complex and lead to the translational repression of the non-optimal mRNAs (**Fig. 7G**).

## Discussion

Our findings reveal the potent and pervasive roles of ZC3H7A and ZC3H7B, two paralog proteins in Chordates, in codon-dependent regulation of mRNA translation and decay. Based on these finding we propose a bifurcated mechanism through which, upon engagement with paused ribosomes on non-optimal mRNAs, the ZC3H7A/B proteins suppress new rounds of translation initiation via recruitment of GIGYF2/4EHP and promote mRNA degradation via recruitment of the CCR4-NOT complex (**Fig. 7G**). In parallel, direct binding of the ZC3H7A/B proteins to the mRNA 3’ UTR leads to CCR4-NOT mediated degradation of the target mRNAs (**Fig. 6F**).

Translation initiation on non-optimal mRNAs is repressed [22, 45, 46], likely due to reduced binding of eIF4E to the 5’ cap [22]. This may represent a feedback mechanism to alleviate the load of elongating ribosomes on non-optimal mRNAs, ultimately enhancing protein synthesis efficiency. Notably, this phenomenon appears to be independent of CNOT3 or DDX6 and could not be attributed to any known sensors of non-optimal codon usage [22]. Crucially, high-throughput and structural biology analyses of the role of CNOT3 in codon-dependent mRNA stability revealed that in human cells, unlike in yeast, it is not the nucleotide type at the wobble site (A/U3 vs. G/C3) but rather the presence of a specific sequence in the D-arm of certain arginine tRNAs at the P-site of the paused ribosome that triggers CNOT3-mediated mRNA decay [21]. Furthermore, high-throughput analysis of mRNA expression in CNOT3-depleted human acute myeloid leukemia, revealed overexpression of the G/C3-rich mRNAs along with downregulation of the A/U3-rich mRNAs [23]. These findings dispute a role for CNOT3 in mediating the repression of A/U3-rich non-optimal mRNAs in humans and suggest that factors other than CNOT3 may interact with ribosomes paused at these non-optimal sequences and trigger translational repression and mRNA decay. Thus, the present study addresses a critical gap in our understanding of the mechanism of codon-dependent post-transcriptional regulation of gene expression in higher eukaryotes by demonstrating the pervasive impacts of ZC3H7A/B on A/U3-rich non-optimal mRNAs.

While the majority of ZC3H7A/B PAR-CLIP reads map to mRNAs, significant proportions (17.5% for ZC3H7A and 5% for ZC3H7B) originate from tRNAs (**Fig. 1C**). Notably, over 50% of the tRNA binding sites for both ZC3H7A and ZC3H7B mapped to tRNA-Arg-AGA (UCU anticodon; **Supp. Table 1**). The presence of specific arginine codons in the ribosomal P-site has been shown to serve as a strong signal for the recruitment of CNOT3 [21]. However, this effect was selective for specific arginine codons (CGG, CGA, and AGG) and did not extend to other arginine codons (AGA, CGC, and CGU). This raises the intriguing possibility that ZC3H7A/B may sense a distinct set of codons compared to CNOT3. Comparative studies exploring the effects of ZC3H7A/B and CNOT3 depletion on ribosomal pausing at various codons -and the subsequent impacts on mRNA translation and stability-will be of significant interest.

The translation-dependent decay process is highly conserved across eukaryotic species and has even been observed in bacteria [14]. However, there are significant differences in codon optimality across species, and factors implicated in *codon optimality-mediated mRNA degradation* in one species (e.g., DDX6 and CNOT3) may not necessarily perform orthologous functions in other species [14, 21]. The ZC3H7A/B genes emerged in Chordates and have no known orthologs in lower eukaryotes. This suggests that higher eukaryotes may possess additional layers of complexity in the regulation of *codon optimality-mediated mRNA degradation* that cannot be fully elucidated through studies of lower eukaryotic models such as yeast, worm, and fly.

This study ascribes a role for the GIGYF2/4EHP translation repressor complex in ZC3H7A/B-mediated inhibition of translation initiation on non-optimal mRNAs. Specifically, upon engagement with paused ribosomes on non-optimal mRNAs, ZC3H7A/B proteins may recruit the GIGYF2/4EHP complex to displace eIF4E from the cap, thereby reducing translation initiation rates. Conspicuously, a similar role for GIGYF2/4EHP has been demonstrated in repressing translation initiation following the detection of ribosome collisions by ZNF598 and EDF1, which interact with the GIGYF2/4EHP complex [42–44]. In addition to GIGYF2/4EHP, ZC3H7A/B proteins also interact with the CCR4-NOT complex, which promotes the ZC3H7A/B-mediated mRNA degradation (**Fig. 6**). The factors that determine whether ZC3H7A/B prioritize translational repression or degradation of non-optimal mRNAs remain unclear. Our data suggest that the interaction between ZC3H7A/B and CNOT1 as well as GIGYF2/4EHP -and consequently the effect on mRNA decay and translation initiation-is independent of DDX6 and CNOT3. However, DDX6 and CNOT3 both interact with ZC3H7A/B, although the functional significances of these interactions are unknown. Future studies will be necessary to delineate the precise mechanisms by which these factors interact and influence each other to achieve coordinated regulation of the translation and stability of non-optimal mRNAs.

Our data indicate that, in addition to binding within the CDS, ZC3H7A/B also bind cellular mRNAs in their 3’ UTRs, leading to CCR4-NOT-mediated decay. Interestingly, we observed a negative correlation between CDS G/C3 content and ZC3H7A/B binding to 3’ UTRs, albeit less prominently than the correlation observed for binding within the CDS (**Supp. Fig. 4A-D**). The mechanism by which CDS codon content could influence ZC3H7A/B binding to 3’ UTRs remains unclear. Previous studies have reported a negative correlation between CDS G/C content, and to a lesser extent 3’ UTR G/C content, and localization to P-body granules [11]. Codon optimality can also affect the microRNA targeting efficacy within the 3’ UTR [47]. Further functional validation is necessary to determine whether CDS codon content directly influences ZC3H7A/B binding to the 3’ UTR or affects downstream mechanisms involved in repressing target mRNA via CCR4-NOT following ZC3H7A/B binding to the 3’ UTR.

Our PAR-CLIP (**Fig. 1I**), RNA-Seq, and Ribo-Seq assays (**Fig. 4E & F**) revealed a high degree of overlap between the mRNA targets of ZC3H7A and ZC3H7B proteins. Additionally, ZC3H7A and ZC3H7B exhibited highly similar patterns of interaction with other proteins, including ribosomes (**Fig. 7B**), components of the CCR4-NOT complex (**Fig. 6E**), and the GIGYF2/4EHP complex (**Fig. 7D & Supp. Fig. 11C**). These findings suggest that ZC3H7A/B proteins have at least partially redundant functions, a common phenomenon for proteins encoded by paralog genes [48]. These redundant functions may also complicate the efforts to define their roles in biological processes using high-throughput loss-of-function approaches such as CRISPR screens, leading to false-negative data. However, despite these redundancies, individual depletion of ZC3H7A or ZC3H7B has, in some cases, produced distinct phenotypes. For instance, in glioblastoma stem cells, CRISPR-mediated depletion of ZC3H7A increased sensitivity to Temozolomide [27]. Conversely, a separate study reported that CRISPR-mediated depletion of ZC3H7B increased the tumorigenicity of familial glioma cells [49]. These distinct phenotypes upon depletion of individual paralogs may arise from cell-type-specific differences in ZC3H7A and ZC3H7B expression or functions.

We observed a stronger correlation between changes in mRNA expression and codon usage in ZC3H7A-or ZC3H7B-overexpressing cells (**Fig. 4A & B**) compared with ZC3H7A/B-double depletion cells (**Fig. 5K & L**). This may be attributed to stochastic changes and adaptation in gene expression during the multi-week selection process required for CRISPR-mediated knockout of ZC3H7A followed by shRNA-mediated knockdown of ZC3H7B in the double depletion cells, which could result in reduced sensitivity to ZC3H7A/B depletion. In contrast, doxycycline-induced overexpression of ZC3H7A or ZC3H7B occurred over a much shorter timeframe (24 hours), likely minimizing adaptation and providing a more accurate representation of the effects. Alternatively, it is possible that ZC3H7A and ZC3H7B are present at sub-stoichiometric levels relative to paused ribosomes in HEK293 cells, and their overexpression significantly amplifies the adverse effects of ribosome pausing on mRNA stability and translation. A similar phenomenon has been reported for ZNF598, a key sensor of collided ribosomes [50, 51]. However, we cannot rule out the possibility that ZC3H7A and ZC3H7B overexpression artificially altered stoichiometric protein levels, leading to exaggerated outcomes.

### Limitations of the study

Our co-IP data demonstrated that ZC3H7A/B proteins bind 80S ribosomes but not eEF1A and eEF2 (**Fig. 7**), two abundant elongation factors typically found on actively translating ribosomes [52]. Notably, besides mRNAs and tRNAs, our PAR-CLIP data (**Supp. Table 1**) also suggest direct interactions between the 5S rRNA and ZC3H7A/B proteins. 5S rRNA is implicated in relaying information regarding the status of the P-site and the A-site to the ribosomal decoding center [53]. Additionally, we demonstrate that the ZC3H7A/B proteins interact with DDX6 and CNOT3-effectors of codon optimality-mediated mRNA degradation, the latter of which occupies the vacant ribosomal E site in the absence of eEF1A and eEF2 [14]. Importantly, all co-IP assays were conducted in the presence of ribonuclease to eliminate potential RNA-bridging artifacts. Based on these interaction patterns and the strong preference of ZC3H7A/B proteins for binding to non-optimal A/U3 mRNAs (**Fig. 2**), it is conceivable that ZC3H7A/B interact with ribosomes paused at non-optimal codons. However, a limitation of this study is the lack of structural information on the interactions between ZC3H7A/B and the ribosome. Additionally, it is unclear whether ZC3H7A/B interact with the paused ribosomes and their associated factors (e.g. CNOT3 and DDX6) directly or through other, yet unknown, intermediary proteins. Our data further show that ZC3H7A/B association with the paused ribosomes likely occurs independently of CNOT3 or DDX6, as these interactions persist in cells knocked down for either CNOT3 or DDX6 (**Fig. 7C**). Nevertheless, we cannot exclude the possibility that residual levels of CNOT3 and DDX6 are sufficient to mediate bridging between ZC3H7A/B and the paused ribosomes. Furthermore, CNOT3 and DDX6 may be needed for efficient ZC3H7A/B-facilitated degradation of non-optimal mRNAs, even if they are not required for the interactions between ZC3H7A/B and paused ribosomes.

## Material and methods

### Cell lines and culture conditions

HEK293 cells (Thermo Fisher Scientific) were cultured in DMEM (Dulbecco’s Modified Eagle Medium; Gibco, Cat. # 41965039) supplemented with 10% Fetal Bovine Serum (FBS; Gibco, Cat. # 10270106) and 100 U/mL penicillin, 100 µg/mL streptomycin (Gibco, Cat. # 15070063). Flp-In T-REx 293 cells (Thermo Fisher Scientific, Cat. # R78007) were grown in high glucose DMEM supplemented with 10% v/v FBS, 100 U/mL penicillin, 100 µg/mL streptomycin, 100 µg/mL zeocin (Invitrogen, Cat. # 460509) and 15 µg/mL blasticidin (BioShop, Cat. # BLA477). Cell lines expressing inducible Flag/HA-ZC3H7A or Flag/HA-ZC3H7B were generated as described previously [54] and selected and maintained in media supplemented with 100 µg/mL hygromycin (BioShop, Cat. # HYG002.1). Expression of tagged proteins was induced for 24 h by the addition of doxycycline (BioShop, Cat. # DOX444) to a final concentration of 1 µg/mL. In this manuscript the cells expressing Flag/HA-Empty are labeled as CTRL and those that express the Flag/HA-tagged protein as OE (i.e. ZC3H7A-OE and ZC3H7B-OE). The Flp-In T-REx HEK293 cells are labeled as Parental and the CRISPR-Cas9-mediated knockout as KO. In all experiments KO cells were compared with their Parental cells, whereas OE cells were compared to CTRL cells. For experiments in the rescued cell line, cells expressing full-length were established as described previously [54] using the relevant KO cells. The cells were selected and maintained in media supplemented with 100 μg/mL hygromycin. All cells were cultured at 37℃, in a humidified atmosphere with 5% CO_2_ and regularly tested for presence of mycoplasma contamination using mycoplasma detection kit (abm, Cat. # G238). ZC3H7A and ZC3H7B double-depletion cells were generated using shRNA to knockdown ZC3H7B in ZC3H7A-KO HEK293 cells. As a control, the Parental cells were transduced with non-targeting control shRNA.

### Generation of knockout cell lines by CRISPR-Cas9

CRISPR-Cas9-mediated genome editing of Flp-In T-REx HEK293 cells was performed as previously described [55]. The oligodeoxynucleotides designed to encode sgRNAs targeting the coding region of the gene of interest are detailed in Supp. Table 4. In brief, the forward and reverse strand oligodeoxynucleotides were annealed and then ligated into the linearized pSpCas9(BB)-2A-GFP plasmid (PX458) (Addgene, plasmid 48138) using BbsI (Thermo Fisher Scientific, Cat. # ER1011). Following transformation, plasmids containing the guide sequence were isolated and verified by sequencing. To create the gene knockout cell lines, 1.3 x10^5^ cells were transfected with the corresponding guide sequence-containing pSpCas9(BB)-2A-GFP plasmid. 24 h post-transfection, GFP-positive cells were single-cell sorted by FACS (BD Aria III Cytometer, Serial. # P18400035) into two 96-well plates and cultivated until colonies formed. Clonal cell lines were assessed by western blotting for the absence of the protein and further analyzed for indel mutations in the targeted alleles using PCR. The PCR products were cloned using the Zero Blunt PCR Cloning Kit (Thermo Fisher Scientific, Cat. # K270040), and ten clones per cell line were sequenced.

### shRNA-mediated gene knockdown

The following lentiviral vector delivered shRNAs were used for generation of stable knockdown cells in this study: Non-Targeting shRNA Controls (Sigma, Cat. # SHC002) and shRNA against human ZC3H7B (Sigma, Cat. # TRCN0000 156115). For packaging, 8 x10^5^ HEK293T cells were cultured in a 6 well plate in high glucose DMEM supplemented with 10% v/v FBS for 24 h. Medium was replaced with OptiMEM (Thermo Fisher Scientific, Cat. # 31985047) 30 min before transfection. Lentivirus particles were produced by transfecting the cells using Lipofectamine 2000 (Thermo Fisher Scientific, Cat. # 11668019) and 1.5 µg lenti-shRNA plasmid, 1 µg psPAX2 (Addgene, plasmid 12260) and 0.5 µg pMD2.G (Addgene, plasmid 12259) packaging plasmids. 6 h post-transfection, the medium was replaced with fresh high glucose DMEM supplemented with 10% v/v FBS. Supernatant was collected at 48 h post-transfection, replaced with 2 mL fresh medium and harvested again after 24 h. Viral particles were briefly centrifuged at 1,500 x g for 5 min at room temperature and purified by filtration (Filtropur S, 0.2 µm, Sarstedt, Cat. # 83.1826.001). Virus solution was stored at −80°C without cryopreservative in 1 mL aliquots or used to infect the cells directly, followed by selection of the stable cell lines with 5 µg/mL puromycin (Merck. Cat. # P8833).

### siRNA-mediated gene knockdown

The following siRNAs were used in this study: ON-TARGETplus Non-targeting Control Pool (Dharmacon, Cat. # D-001810-10-50), CNOT1 siRNA SMARTpool (Dharmacon, Cat. # L-015369-01-0005), CNOT3 siRNA SMARTpool (Dharmacon, Cat. # L-020319-00-0005), DDX6 siRNA SMARTpool (Dharmacon, Cat. # L-006371-00-0005). Cells were reverse transfected using Lipofectamine RNAiMAX (Thermo Fisher Scientific, Cat. # 13778075) and corresponding siRNA at the final concentration of 10 pmol and cultured in high glucose DMEM supplemented with 10% v/v FBS for 48 h. The transfection media was replaced with high glucose DMEM supplemented with 10% v/v FBS, 100 U/mL penicillin, and 100 µg/mL streptomycin, and cells were harvested after another 24 h incubation.

### Plasmid cloning

pENTR4 plasmids for cloning of Flag/HA-tagged proteins in Flp-In T-REx HEK293 cells were generated by PCR amplification of the respective coding sequences of the gene of interest (GOI) from cDNA of Flp-In T-REx HEK293 cells using primers flanked by attB1 and attB2 recombination. PCR products were recombined into the pDONR221 plasmid (Thermo Fisher Scientific, Cat. # 12536017) using the GATEWAY BP system (Thermo Fisher Scientific, Cat. # 11789020). The resulting pENTR4-GOI plasmids were recombined with the pFRT/TO/FH-DEST destination vector using the GATEWAY LR recombinase system (Thermo Fisher Scientific, Cat. # 11791020). To generate the plasmids encoding λN-v5-ZC3H7A or λN-v5-ZC3H7B, full length and truncated sequences were PCR-amplified using specific primers (**Supp. Table 4**) and cloned into the pCI-neo Lambda N-v5 plasmid (Promega, Cat. # E1841) using NotI (Thermo Fisher Scientific, Cat. # FD0593) and EcoRI (Thermo Fisher Scientific, Cat. # FD0274) restriction enzymes. Chemically competent *E. coli* DH5α cells (Thermo Fisher Scientific, Cat. # 18265017) were used for plasmid amplification. The final sequences were verified using Sanger sequencing.

### PAR-CLIP

24 h after induction of expression of Flag/HA-ZC3H7A or Flag/HA-ZC3H7B with doxycycline (1 μg/mL) PAR-CLIP was performed on whole cell lysates as described previously[56]. The resulting PAR-CLIP cDNA libraries were sequenced on an Illumina HiSeq2500 instrument, and the data were analyzed using the PAR-CLIP suite [56] and PARalyzer [57, 58]. For metagene analysis, reads mapping to mRNAs or mRNA precursors with d1 T-to-C sequence transitions and a length of ≥20 nt were extracted and aligned to the human genome using STAR version 2.5.2a [59]. For the metagene analysis of reads mapping to different mRNA regions, gencode gtf file (gencode.v19.chr_patch_hapl_scaff.annotation.gtf) was used to calculate the logarithm of average read coverage for 3’ UTR, CDS, and 5’ UTR for the 500 mRNAs with highest coverage. Relative UTR and CDS sizes were calculated based in their average size in all mRNAs expressed in HEK293. Average coverage was calculated for each UTR and CDS independently according to the actual length of the region and number of bins (10 for 5’ UTR, 60 for CDS and 20 for 3’ UTR) and additionally represented in the upper panel as percent of total gene coverage. The MetaPlotR Perl/R pipeline [60] was used for plotting metagenes. *De novo* motif enrichment analysis was performed using the MEME suite, specifically DREME (http://meme-suite.org/tools/meme). The analysis focused on the top 500 clusters in the CDS and 3’ UTR regions, as defined by PARalyzer, with non-target mRNAs serving as controls.

### Polysome profiling

Cells were cultures to a maximum 80% confluency and washed with PBS containing 100 μg/mL Cycloheximide (Sigma, Cat. # 01810), collected via scraping and pelleted by centrifugation at 4°C for 5 min. The pelleted cells were lysed in 500 μl buffer (5 mM Tris-HCl, pH 7.5, 2.5 mM MgCl2, 1.5 mM KCl, complete) supplemented with 0.5% w/v sodium deoxycholate, 0.5% w/v Triton X-100, 100 μg/mL, Cycloheximide, 2 mM DTT, EDTA-free protease inhibitor cocktail (Roche, Cat. # 04693159001), and 200 U/ mL RiboLock RNase Inhibitor (Thermo Fisher Scientific, Cat # EO0382). The lysates were cleared by centrifugation at 20,000 × g for 5 min at 4°C and the RNA concentration was measured by NanoDrop 2000 (Thermo Fisher Scientific). Equivalent of 400 µg of RNA was loaded onto 10%–50% sucrose gradients and centrifuged at 36,000 x rpm for 2 h at 4°C using SW40Ti rotor in Optima L-80XP ultracentrifuge (Beckman Coulter). UV absorbance (254 nm) was using an ISCO gradient fractionation system, and the optical density was continuously recorded with a Foxy JR Fractionator (Teledyne ISCO).

### Ribo-Seq (ribosome profiling) and RNA-Seq

Flag/HA control, ZC3H7A-OE, or ZC3H7B-OE HEK293 cells (3 independent replicates each) were resuspended in lysis buffer (5 mM Tris-HCl (pH 7.5), 2.5 mM MgCL2, 1.5 mM KCl, 100 μg/mL Cycloheximide, 1x EDTA-free protease inhibitor cocktail, 2 mM DTT, 200 U/ml RiboLock RNase Inhibitor, 0.5% (w/v) Triton X-100, and 0.5% (w/v) Sodium Deoxycholate), to isolate the polysomes. A 50 µg fraction of each lysate was collected for polyA RNA-Seq and 500 µg of the lysates were subjected to ribosome footprinting by RNase I (Invitrogen, Cat. # AM2294) treatment at 4°C for 60 min with end-over-end rotation. Ribosomes were pelleted by ultracentrifugation in a 34% sucrose cushion at 70,000 rpm for 3 h and RNA fragments were extracted twice with acid phenol, once with chloroform, and precipitated with isopropanol in the presence of NaOAc and GlycoBlue (Thermo Fisher Scientific, Cat. # AM9515). Purified RNA was resolved on a denaturing 15% polyacrylamide-urea gel and the section corresponding to 28-32 nucleotides containing the RPFs was excised, eluted, and precipitated by isopropanol. All samples were analyzed on a Bioanalyzer small RNA chip (Agilent Technologies) to confirm expected size range and quantity, followed by rRNA depletion using the Illumina Ribo-Zero rRNA Depletion Kit, PNK end repair, generation of libraries with NEXTFLEX® Small RNA-Seq Kit v3 (BIOO Scientific, Cat. # 5132-05), and sequencing on Illumina HiSeq4000 platform with 50 nt single-end reads to depths of 100 million read per sample, and data files were provided in fastq format. Raw fastq files were trimmed using cutadapt, following BIOO Scientific recommendations by trimming 3’ adapter and 4 bases from either side of each read. As a filtering step, trimmed reads were first aligned to non-coding RNAs (http://ftp.ensembl.org/pub/release-103/fasta/homo_sapiens/ncrna/Homo_sapiens.GRCh38.ncrna.fa.gz) using bowtie v1.1.0 before remaining unaligned reads were aligned to the human genome (hg38) using STAR [61]. Aligned reads were used as input to estimate the quality of libraries including estimation of P-site, visualisation of start/stop windows, metagene analysis, and assessment of reading frame distribution. Counts per gene were calculated using HTSeq-count [62].

Total RNA was extracted from the aliquots described in the paragraph above, using TRIzol reagent (Invitrogen, Cat # 15596018), according to the manufacturer’s instructions. Oligo(dT)-selected RNAs were reverse transcribed into cDNA for RNA-sequencing using the Illumina TruSeq RNA Sample Preparation Kit v2, following the manufacturer’s instructions. Sequencing was performed on an Illumina HiSeq4000 platform with 50 nt single-end reads to depths of 100 million read per sample, and data files were provided in fastq format.

The RNA-Seq and Ribo-Seq analyses of the ZC3H7A and ZC3H7B double-depletion (ZC3H7A/B-DD) and the relevant control cells (3 independent replicates each) were performed by EIRNA Bio (Cork, Ireland). In brief, flash frozen cell pellets were lysed in ice-cold polysome lysis buffer (20 mM Tris pH 7.5, 150 mM NaCl, 5 mM MgCL2, 1 mM DTT, 1% Triton X-100, and 100 µg/mL cycloheximide). For stranded RNA-Seq, total RNA was extracted from 10% of lysate using TRIzol, before being rRNA depleted, fractionated, and converted into Illumina compatible cDNA libraries. For Ribo-Seq, remaining lysates were digested in the presence of 35 U RNase I for 1 h at room temperature. Following RNA purification, PNK end repair and size selection of ribosome protected mRNA fragments on 15% urea PAGE gels, contaminating rRNA was depleted from samples using EIRNA Bio’s custom biotinylated rRNA depletion oligos. Enriched fragments were converted into Illumina compatible cDNA libraries. Both stranded RNA-Seq libraries and Ribo-Seq libraries were sequenced on Illumina’s Nova-seq 6000 platform with 150 pair-end to depths of 20 million and 100 million read pairs per sample respectively.

### mRNA features analysis

The cDNA, CDS, 5’ and 3’ UTRs of the longest expressed transcripts were retrieved from Ensembl BioMart using the biomaRt Bioconductor package [63]. Length and G/C content were calculated using bedtoolsnuc (bedtools v2.25.0) [64]. The length of cDNA, CDS, and UTRs between groups (up-regulated, down-regulated, or unchanged) were compared using Welch’s Two Sample t-test and cumulative distribution plots were generated to visualise the distribution of Log_2_ fold changes. Similarly, genes were grouped into six categories [0 sites, 1−2 sites (A), 3-4 sites (B), 5-9 sites (C), 10-24 sites (D), ≥ 25 sites (E)] based on the number of PAR-CLIP binding sites present within the region of interest. Length and G/C content were compared across these categories using the non-parametric Dunn test, applied with Holm multiple test correction. A p-value of less than 0.05 was considered statistically significant. Codon usage per gene was obtained from DNA Hive (https://dnahive.fda.gov/dna.cgi?cmd=codon_usage&id=537&mode=cocoputs) using the dataset o537-Human_CDS.tsv [65]. Codon usage frequencies were retrieved from the Kazusa Codon Usage Database (https://www.kazusa.or.jp/codon/cgi-bin/showcodon.cgi?species=9606&aa=1&style=N). Codon efficiency per gene/CDS was calculated by multiplying the number of each codon by its respective usage frequency per CDS, summing these values to generate an overall efficiency score per gene. This sum was then divided by the CDS length to normalise efficiency. To assess the correlation between codon usage and differential gene expression, log fold-change (LogFC) per gene was correlated with codon usage per gene, where LogFC values for genes were separately correlated with usage of each of the 64 codons. These correlations were ranked from high to low and visualised as a waterfall plot, with codons ending in A/T(U) coloured red and those ending in G/C coloured blue, following the methodology described in [66].

### RNA extraction and RT-qPCR

Cells were harvested, and RNA was isolated using TRIzol RNA extraction kit. 1 µg purified total RNA was treated with DNase I (Thermo Fisher Scientific, Cat. # EN0521). Total RNA was reverse-transcribed with Superscript III reverse transcriptase (Invitrogen, Cat. # 18080085), with 100 ng random hexamers (Thermo Fisher Scientific, Cat. # N8080127). mRNA abundance was quantified using the LightCycler 480 SYBR Green I Master mix (Roche, Cat. # 04887352001) and LightCycler® 480 real-time PCR system according to the manufacturer’s protocol. All primers are listed in Supp. Table 4.

### *In vitro* mRNA synthesis

Synthetic mRNAs were generated for the CDS of human *KRAS* (CCDS8703.1) and *HRAS* (CCDS7698.1), which were tagged with an in-frame Flag epitope at their C-terminus (*KRAS-FLAG* and *HRAS-FLAG*, respectively). The translationally active 5’ and 3’ UTRs were designed by RIBOPRO (Oss, The Netherlands), who also equipped the mRNAs with a 5’ Cap1 structure and a 150 nucleotide-long poly(A) tail. mRNA quality was assessed by spectrophotometry and gel electrophoresis by the manufacturer (RIBOPRO, The Netherlands).

### Analysis of mRNA stability

HEK293 cells were transfected with 5 ng of the *FL* control plasmid, 10 ng of *RL*-5BoxB reporter and 100 ng of v5-tagged λN-empty, λN-ZC3H7A or λN-ZC3H7B tethering plasmids in a 12 well plate using Lipofectamine 2000. 24 h post-transfection, cells were treated with 15 μg/mL actinomycin D (Thermo Fisher Scientific, Cat. # A9415) to stop the de novo transcription. Cells were collected at desired timepoints, followed by RNA extraction and RT-qPCR as described above. The sequences of the primers are listed in **Supp. Table 4**.

To measure the stability of the synthetic *KRAS-FLAG* and *HRAS-FLAG* mRNAs, 150,000 doxycycline inducible Flag/HA control or ZC3H7B-OE HEK293 cells were cultured in 6 well plate. Expression of Flag/HA-ZC3H7B was induced for 24 h by the addition of doxycycline (1 μg/mL), followed by transfection with 500 ng of *KRAS-FLAG* or *HRAS-FLAG* mRNA using Lipofectamine 2000. After 5 h incubation, the transfection media was replaced with high glucose DMEM supplemented with 10% v/v FBS, 100 U/mL Penicillin, 100 µg/mL streptomycin and the samples were collected at the indicated timepoints. RNA extraction and RT-qPCR was done as described above. The sequences of the primers are listed in **Supp. Table 4**.

### RNA immunoprecipitation (RNA-IP)

15 x10^6^ Dox-inducible Flag/HA-ZC3H7B HEK293 cells were cultured in 15 cm plates. The expression of tagged proteins was induced for 24 h by the addition of doxycycline to a final concentration of 1 μg/mL. Cells were lysed in lysis buffer A containing 40 mM HEPES-KOH; pH 7.5, 120 mM NaCl, 1 mM EDTA, 0.3% CHAPS, 100 U/mL RiboLock RNase inhibitor, and supplemented with complete EDTA-free protease inhibitor (Roche, Cat. # 04693124001) and phosphatase inhibitor (Roche, Cat. # 4906845001). The lysates were centrifuged at 15,000 g for 15 min at 4°C and precleared with 70 μl blocked and pre-washed Dynabeads™ Protein G (Thermo Fisher Scientific, Cat. # 10004D) for 1.5 h at 4°C with gentle agitation. Protein concentration was measured by Bradford assay (Bio-Rad, Cat. # 5000006), and 2 mg of lysates were incubated with 3 μg of anti-Flag antibody for 30 min at 4°C. After adding the blocked and pre-washed Dynabeads™ Protein G, the mixture was incubated overnight at 4°C with gentle rotation. The precipitated beads were then washed three times with 1 mL of lysis buffer A, followed by two washes with wash buffer consisting of 15 mM HEPES-KOH (pH 7.5), 100 mM KCl, 7.5 mM MgCL2, 2 mM DTT, 1% Triton X-100, 100 U/mL RiboLock RNase inhibitor, supplemented with a complete EDTA-free protease inhibitor tablet and resuspended in 1 mL of wash buffer. 10% of the volume was used for western blotting and the remainder was used for RNA extraction using TRIzol, as described above.

### Co-immunoprecipitation (co-IP)

Cells were washed with cold PBS and collected by scraping in the lysis buffer (40 mM HEPES pH 7.5, 120 mM NaCl, 1 mM EDTA, 0.3% CHAPS, supplemented with complete EDTA-free protease inhibitor tablet and phosphatase inhibitor cocktail). Dynabeads™ Protein G were blocked with 2% BSA in PBST for 1 h and washed 3x for 3 min each in TBST. 15 mg of the pre-cleared lysates for Flag/HA-ZC3H7A and 2 mg of the pre-cleared lysates for Flag/HA-ZC3H7B HEK293 cells were incubated with 3 µg anti-Flag antibody and 60 µl of blocked and prewashed Dynabeads™ Protein G at 4°C overnight, in presence of RNase A (Thermo Fisher Scientific, Cat. # EN0531) to eliminate potential RNA bridges. Beads were washed 3× for 10 min each with wash buffer (50 mM HEPES pH 7.5, 150 mM NaCl, 1 mM EDTA, 0.3% CHAPS, supplemented with a complete EDTA-free protease inhibitor tablet and phosphatase inhibitor cocktail) and protein was eluted in SDS loading buffer (0.25% Bromophenol blue, 0.5 M DTT, 50% Glycerol, 10% SDS, and 0.25 M Tris-Cl pH 6.8).

### Western blotting

Cells were washed with ice-cold PBS and detached from the plates using plastic scrapers. After pelleting by centrifugation at 12,000 g for 1 min at 4℃, cell pellets were lysed in RIPA buffer (50 mM Tris-HCL pH 7.4, 150 mM NaCl, 2 mM EDTA, 1% NP-40, 0.1% SDS) supplemented with protease inhibitor (Roche, Cat. # 11836170001). The protein concentration was measured using Bradford protein assay (Bio-Rad, Cat. # 5000006) and 40 µg of total protein was pre-mixed with loading buffer (0.25% Bromophenol blue, 0.5 M DTT, 50% Glycerol, 10% SDS, and 0.25 M Tris-Cl pH 6.8) before boiling at 95℃ for 3 min, followed by a short incubation on ice. Samples were loaded onto an SDS-PAGE gel and wet transferred onto a PVDF membrane (Merck, Cat. # IPFL00010). Thereafter, membranes were blocked with 5% BSA (Thermo Fisher Scientific, Cat. # A9647) at room temperature for 1 h, followed by incubation with the indicated primary antibodies overnight at 4℃ and secondary antibody for 1 h at room temperature. All blots were scanned, and images were captured using the dyssey system (LI-COR, Cat. # ODY-1540) or G:Box chemi XX6 imaging systems (SYNGENE) using SuperSignal™ West Pico PLUS Chemiluminescent Substrate (Thermo Fisher Scientific, Cat. # 34580). The list of antibodies used in this study is provided in **Supp. Table 5**. The uncropped images of all blots are provided in **Supp. Fig. 12-21**.

### Dual luciferase reporter tether-function assay

For the tether-function assay, the λN-BoxB tethering approach [35] was used. Briefly, a *Renilla luciferase (RL)* reporter containing five BoxB hairpins in its 3′ UTR was co-transfected along with the construct encoding a fusion of ZC3H7A or ZC3H7B to a λN peptide, which allows the protein to bind to the reporter’s BoxB elements. The firefly luciferase (FL) expression plasmid was used as control. 150,000 HEK293 cells were transfected in a 24-well plate with 5 ng of the FL plasmid, 20 ng of RL-5BoxB and 100 ng of v5-tagged λN-empty, λN-ZC3H7A or λN-ZC3H7B using Lipofectamine 2000. 24 h after transfection, cells were lysed and luciferase activity was measured with the Dual-Luciferase Reporter Assay (Promega, Cat. # E1960) according to the manufacturer’s instructions using the BMG labtech FLUOstar plate reader. *RL* values were normalized against *FL* levels, and *RL*:*FL* values for the λN-ZC3H7A or λN-ZC3H7B were normalized relative to λN-empty level for each condition.

### mCherry/GFP codon optimality reporter assay

HEK293 cells with the indicated genotypes were seeded at 150,000 cells per well in 24-well plates in complete medium. The cells were then transfected with 250 ng of either the 100% G/C3 or 25% G/C3 mCherry and 250 ng of GFP control plasmids [34] using Lipofectamine 2000. 24 h after transfection, cells were detached and analysed via the BD Accuri C6 flow cytometer; gating recorded 10,000 cells and measured their fluorescence for both GFP and mCherry.

### Cell growth measurement

HEK293 cells with the indicated genotypes were seeded at 50,000 cells per well in 6-well plates in complete medium for the indicated times. Cells were trypsinized and stained with Trypan Blue for 2 min and counted under the microscope using Haemocytometer. The process was repeated for 5 days to monitor the comparative cell growth.

### Sequence alignment and domain analysis

Amino acid sequences were obtained from the NCBI database: ZC3H7A (NCBI Reference Sequence: NP_054872.2); ZC3H7B (NCBI Reference Sequence: NP_060060.3). Protein sequence alignments were carried out using multAlin software (http://multalin.toulouse.inra.fr/multalin/) [67]. Domains analyses were performed via Interpro (http://www.ebi.ac.uk/interpro/). Disorder profile of proteins was mapped via DISOPRED3 [68].

### Statistical Analyses

Statistical tests were performed using R or Prism 6 (GraphPad). Error bars represent standard deviation (SD) from the mean. Number of independent replicates and the statistical analysis used for each assay is described in the relevant figure legends. P values <0.05 were considered significant. Adjusted P value cut-offs are described in the relevant figure legends.

## Acknowledgments

This work was supported by a grant from the Biotechnology and Biological Sciences Research Council (BB/W008165/1) to S.M.J. We thank Dr. Ariel Bazzini (Stower Institute) for providing the mCherry reporters with different percentages of optimal G/C3 codons. We are grateful to Prof RJH Davies for critical reading and editing of the manuscript, Dr. Xu Zhang, Haotian Zhuang and Izzy Shpilman for helpful discussions, and Beth Cunningham, Ciarrai Mcconnell, Dr. Sunghoon Kim, and Dr. Angela Hackett for technical assistance. T.M. is supported by a PhD studentship from Brainwaves Northern Ireland. J.A.W is supported by CRUK core funding (A29252) at the CRUK Scotland Institute.

## Authors contribution

**Conceptualization:** P.H.S., P.N., S.M.J.; **Investigation:** P.H.S., P.N., A.G., S.C., N.J., T.M., S.M.J.; **Methodology:** P.H.S., P.N., A.G., S.M.J.; **Resources:** R.J.K., J.H.C., J.L., S.A.S., X.S.R., C.G.G., N.S., T.T., S.M.J.; **Data curation & analysis:** P.H.S., A.G., J.A.W., S.M., S.M.J.; **Validation:** J.A.W., N.J., S.M.J.; **Visualization:** P.H.S., P.N., A.G., S.M., S.M.J.; **Writing – original draft:** P.H.S., P.N., S.M.J.; **Writing – review & editing:** All authors; **Project administration:** P.H.S., P.N., S.C., S.M.J.; **Funding acquisition:** S.M.J.; **Supervision:** S.M.J.

## Supplementary Tables Legends

**Supplementary Table 1.** PAR-CLIP datasets for analysis of direct interactions between ZC3H7A and ZC3H7B proteins with cellular mRNAs in HEK293 cells.

**Supplementary Table 2.** RNA-Seq and Ribo-Seq dataset for ZC3H7A and ZC3H7B overexpressing HEK293 cells.

**Supplementary Table 3.** RNA-Seq and Ribo-Seq dataset for ZC3H7A/B-double depletion HEK293 cells.

**Supplementary Table 4.** List and sequence of primers and small guide RNAs used in this study.

**Supplementary Table 5:** List of antibodies used in this study.

