## Supplementary Figures for "Codon-dependent regulation of mRNA translation and stability by ZC3H7A and ZC3H7B RNA-binding proteins"

**Supplementary Figures 1-21:**

### Supp. Figure 1; related to Fig. 1

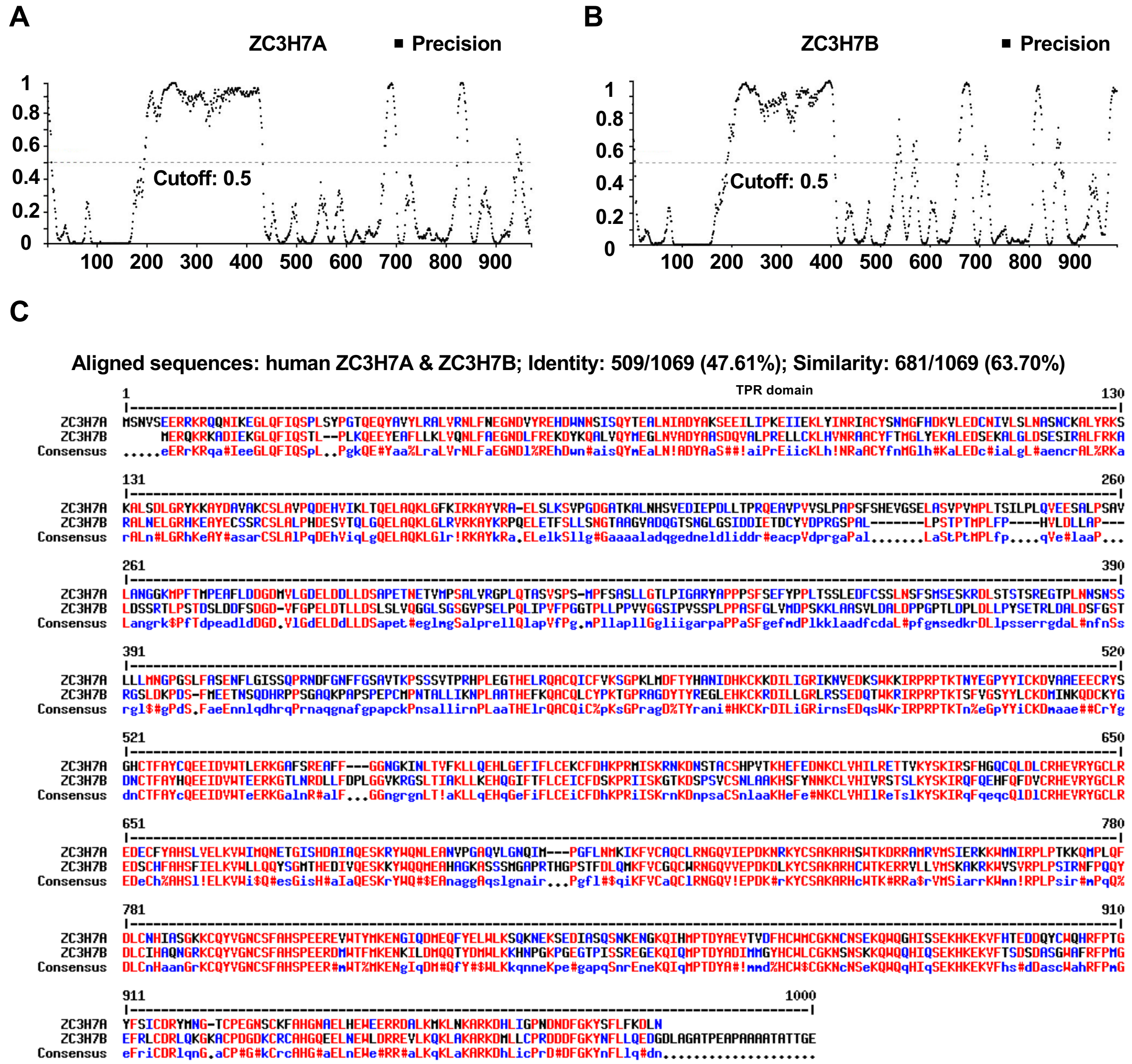


**Supplementary Figure 1. The predicted domains of human ZC3H7A and ZC3H7B proteins; related to Figure 1.** (**A & B**) Disorder profile of human ZC3H7A (NP_054872.2) (A) and ZC3H7B (NP_060060.3) (B) mapped via DISOPRED3. The grey dashed horizontal line marks the threshold above which amino acids are regarded as disordered [1]. (**C**) Alignment of the human ZC3H7A (NP_054872.2) and ZC3H7B (NP_060060.3) protein sequences by multAlin software (http://multalin.toulouse.inra.fr/multalin/multalin.html).

### Supp. Figure 2; related to Fig. 1

A

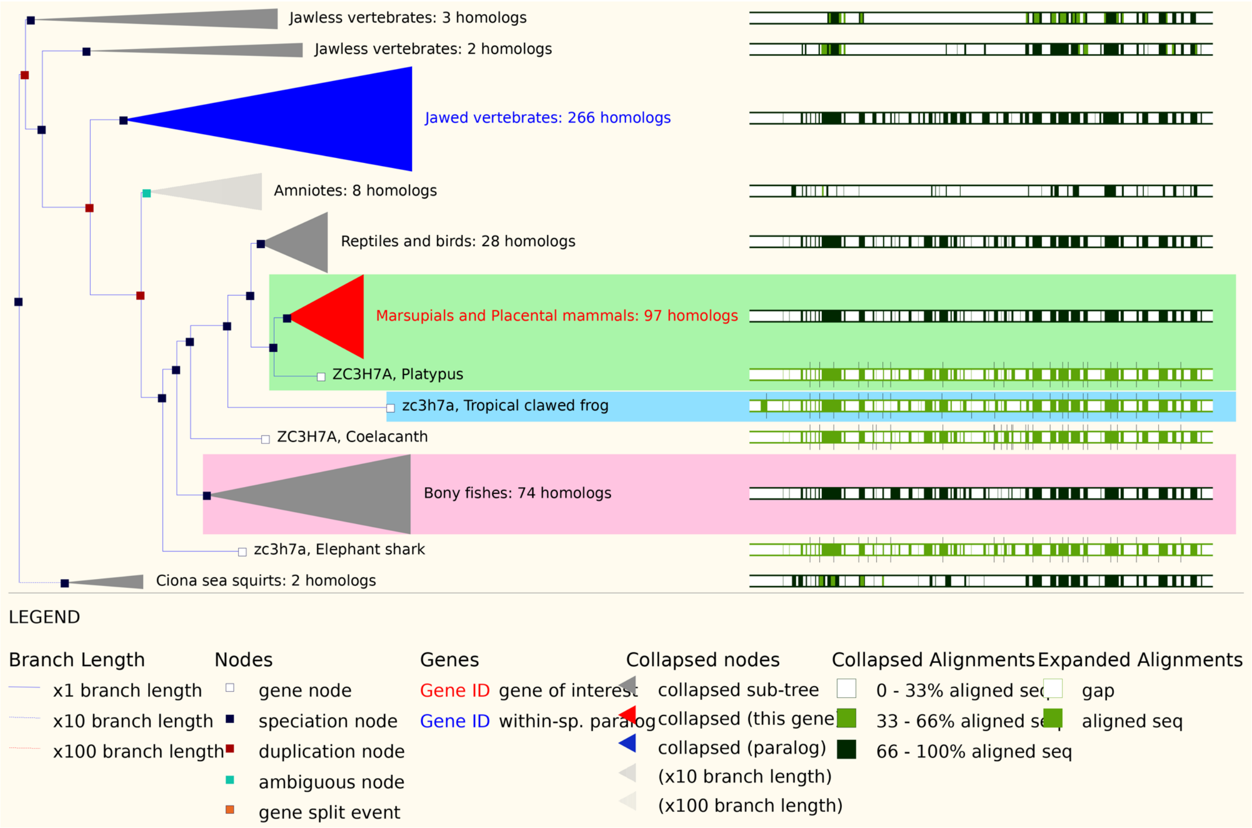


B

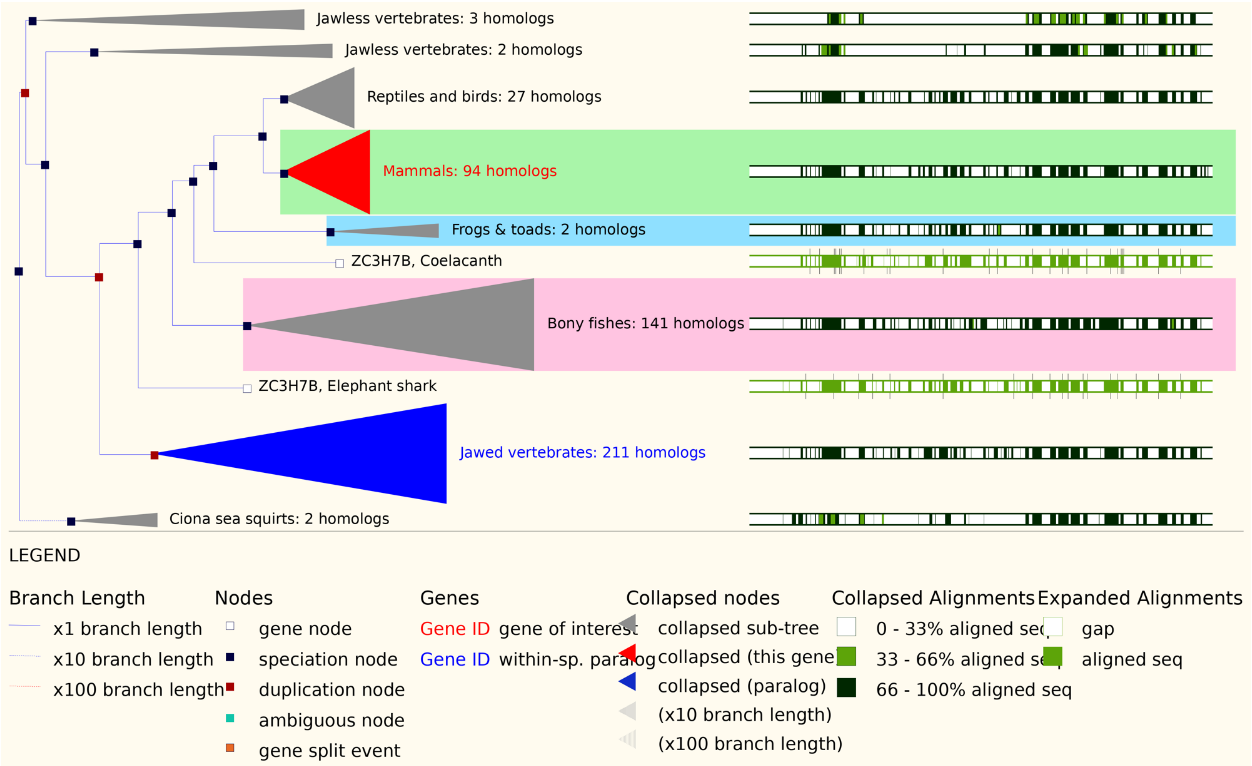


**Supplementary Figure 2. The evolutionary dynamics of the *ZC3H7A* and *ZC3H7B* genes.** (**A & B**) Phylogenetic tree of ZC3H7A (A) and ZC3H7B (B) genes generated by Ensembl Genome Browser (https://www.ensembl.org/index.html).

**Supp. Figure 3; related to Fig. 2**

A

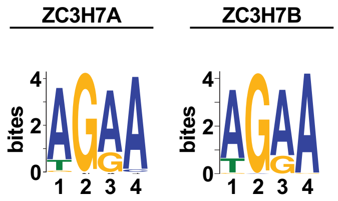


B

| | Key | no. of binding sites | No. of genes: All | CDS | 3′ UTR |
|---|---|---|---|---|---|
| ZC3H7A | black | 0 | 4334 | 6196 | 7073 |
| | grey | 1-2 (A) | 2575 | 2805 | 2444 |
| | blue | 3-4 (B) | 1408 | 1177 | 883 |
| | cyan | 5-9 (C) | 1753 | 1049 | 772 |
| | green | 10-24 (D) | 1316 | 424 | 446 |
| | red | ≥25 (E) | 298 | 33 | 66 |
| ZC3H7B | black | 0 | 3674 | 5901 | 5332 |
| | grey | 1-2 (A) | 1731 | 2478 | 1922 |
| | blue | 3-4 (B) | 1081 | 1225 | 1044 |
| | cyan | 5-9 (C) | 1726 | 1334 | 1439 |
| | green | 10-24 (D) | 2402 | 648 | 1546 |
| | red | ≥25 (E) | 1057 | 85 | 388 |

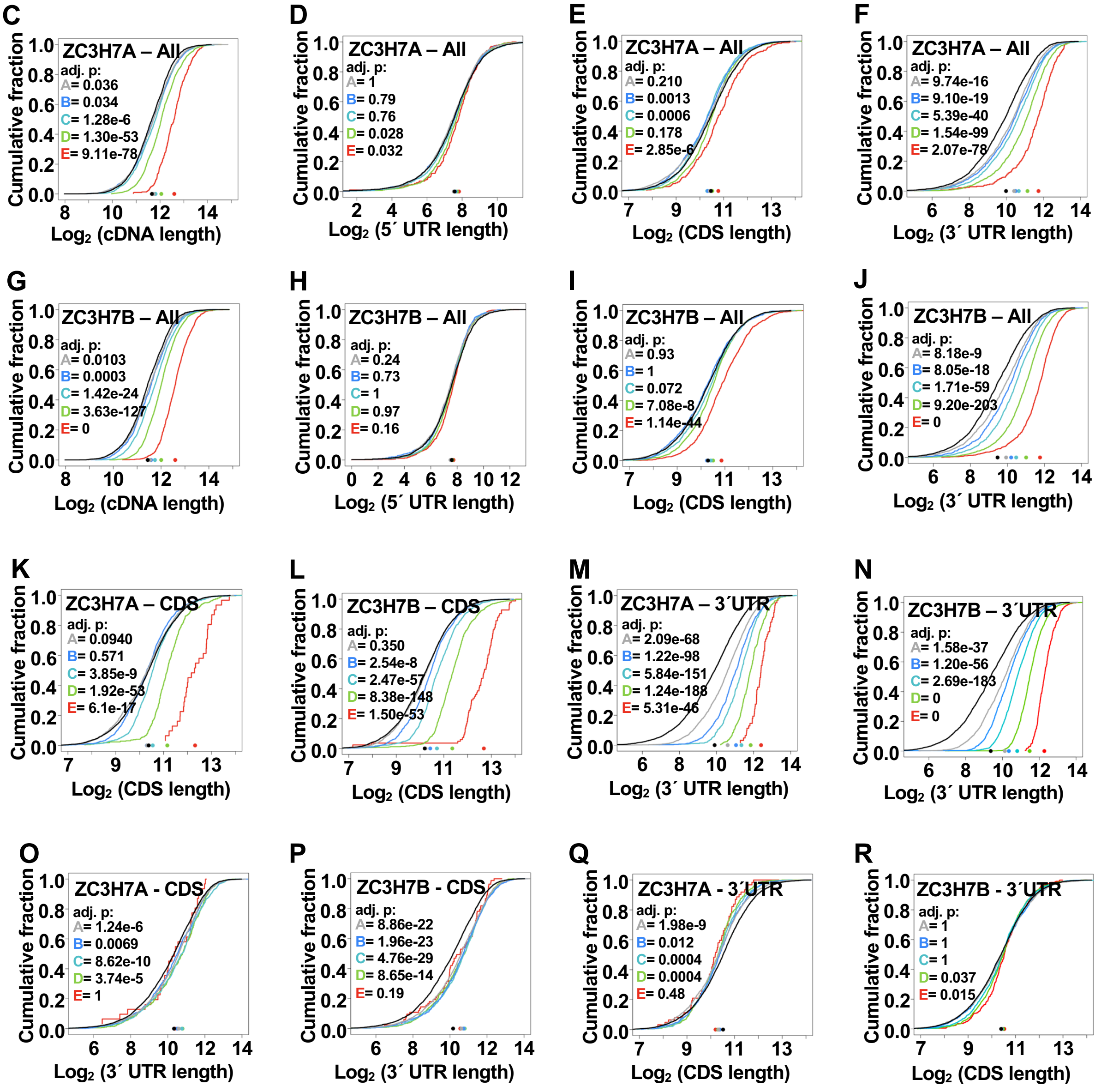


**Supplementary Figure 3. Analyses of sequence features enriched in ZC3H7A and ZC3H7B PAR-CLIP target mRNAs; related to Figure 1.** (**A**) Analysis of sequence motifs enriched in the top 500 PAR-CLIP peaks of ZC3H7A and ZC3H7B. (**B**) Summary of the ZC3H7A or ZC3H7B PAR-CLIP target mRNAs divided based on the number and position of binding sites. (**C-F**) Cumulative distribution analysis of the correlation between the length of the indicated mRNA regions and the number of ZC3H7A binding sites across ZC3H7A target mRNAs. (**G-J**) Cumulative distribution analysis of the correlation between the length of the indicated mRNA regions and the number of ZC3H7B binding sites across ZC3H7B target mRNAs. (**K & L**) Cumulative distribution analysis of the correlation between the CDS length and the number of ZC3H7A (K) or ZC3H7B (L) binding sites across the CDS of ZC3H7A ZC3H7B target mRNAs, respectively. (**M & N**) Cumulative distribution analysis of the correlation between the 3´ UTR length and the number of ZC3H7A (M) or ZC3H7B (N) binding sites across the 3´ UTR of ZC3H7A and ZC3H7B target mRNAs, respectively. (**O & P**) Cumulative distribution analysis of the correlation between the 3´ UTR length and the number of ZC3H7A (O) or ZC3H7B (P) binding sites across the CDS of ZC3H7A and ZC3H7B target mRNAs, respectively. (**Q & R**) Cumulative distribution analysis of the correlation between the CDS length and the number of ZC3H7A (Q) or ZC3H7B (R) binding sites across the 3´ UTR of ZC3H7A and ZC3H7B target mRNAs, respectively.

Supp. Figure 4; related to Fig. 2

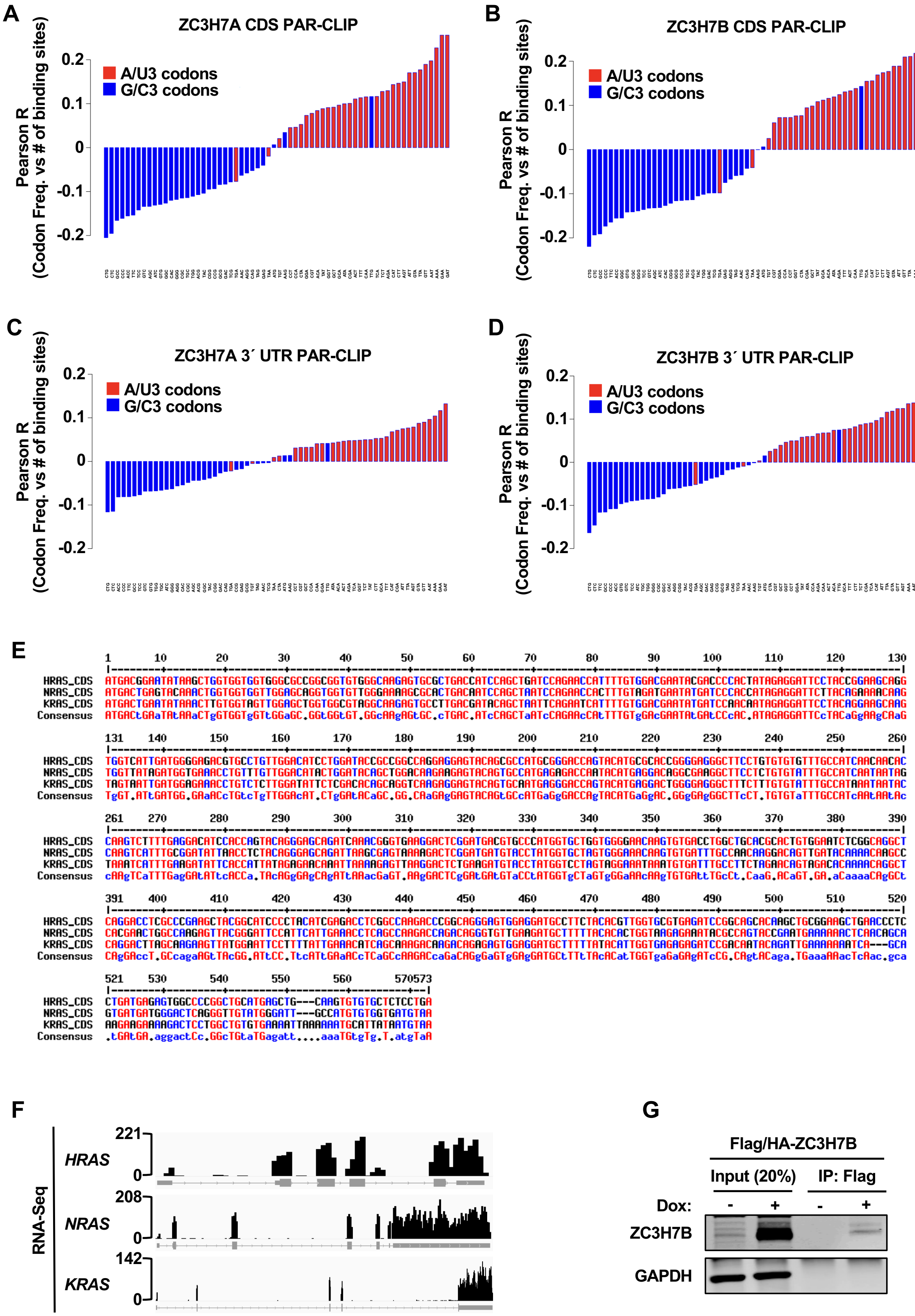


**Supplementary Figure 4. Enrichment for A/U3 non-optimal codons in ZC3H7A/B-bound mRNAs; related to Figure 2.** (**A & B**) Waterfall plots showing Pearson correlations between codon usage frequencies and number of ZC3H7A (A) and ZC3H7B (B) PAR-CLIP binding sites mapped to CDS. (**C & D**) Waterfall plots showing Pearson correlations between codon usage frequencies and number of ZC3H7A (C) and ZC3H7B (D) PAR-CLIP binding sites mapped to 3´ UTR. (**E**) Alignment of the human *HRAS* (CCDS7698.1), *NRAS* (CCDS877.1) and *KRAS* (CCDS8703.1) CDS sequences by multAlin software (http://multalin.toulouse.inra.fr/multalin/multalin.html). (**F**) IGV tracks of the *HRAS*, *NRAS* and *KRAS* mRNAs in HEK293 cells. All three genes are encoded on the (–) strand but are presented in their 5´ to 3´ orientation. (**G**) Western blot analysis with the indicated antibodies in the Flag/HA-ZC3H7B RNA-Immunoprecipitation (RNA-IP) assay.

### Supp. Figure 5; related to Fig. 3

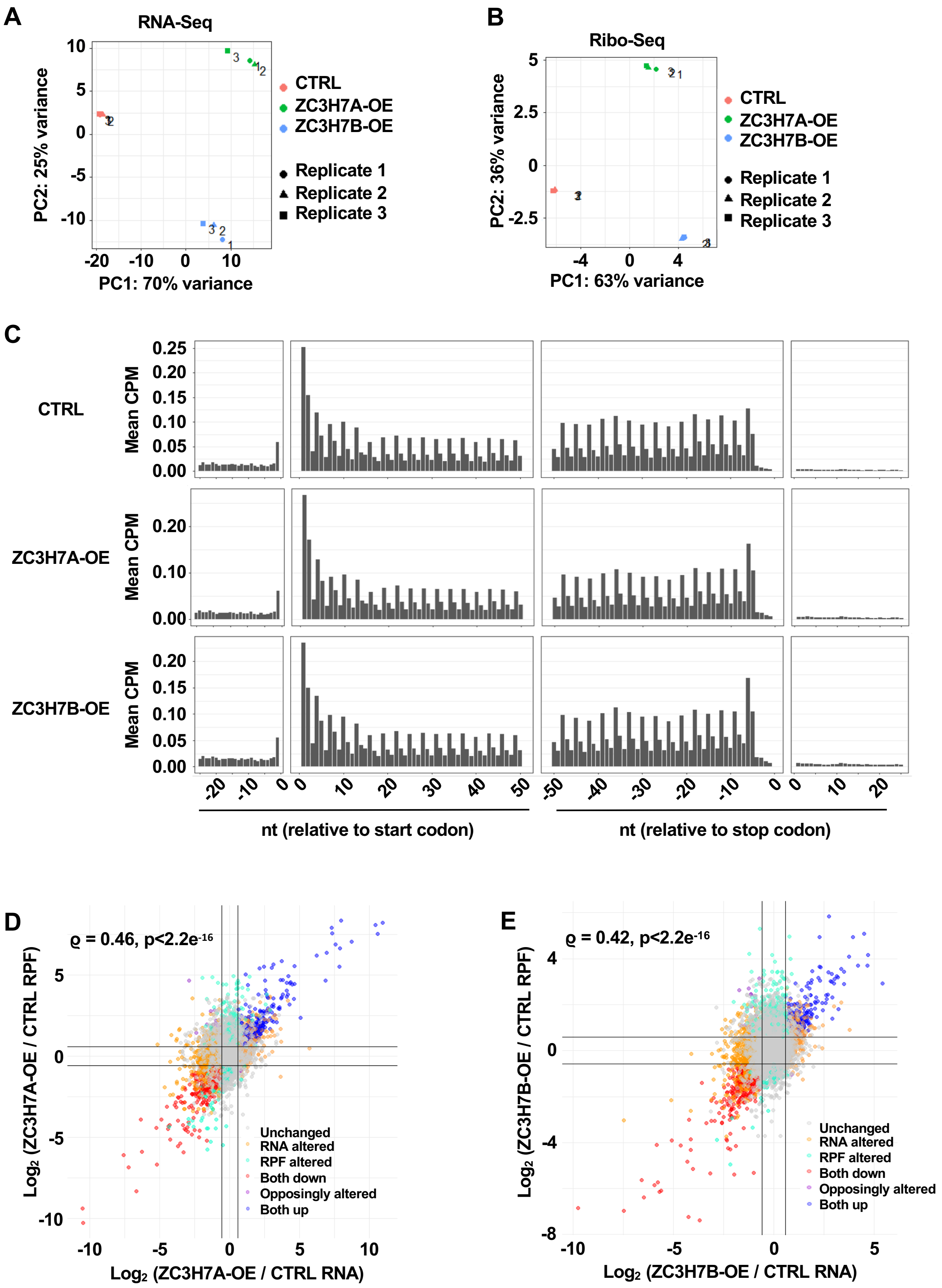


**Supplementary Figure 5. RNA-Seq and Ribo-Seq analyses of Flag/HA-ZC3H7A, Flag/HA-ZC3H7B, and Flag/HA control (CTRL) HEK293 cells; Related to Figure 3.** (**A**) PCA plot of RNA-Seq samples from Flag/HA control, Flag/HA-ZC3H7A, and Flag/HA-ZC3H7B expressing samples. (**B**) PCA plot of RPF (Ribo-Seq) samples from Flag/HA control, Flag/HA-ZC3H7A, and Flag/HA-ZC3H7B expressing samples. (**C**) Frequency of reads for footprint libraries showing the expected

3nt periodicity in relation to the translation start and stop codons in representative replicates from Flag/HA control, Flag/HA-ZC3H7A, and Flag/HA-ZC3H7B expressing samples. (**D & E**) Scatter plot showing the comparison between the changes in mRNA expression vs ribosome occupancy in ZC3H7A-OE (D) and ZC3H7B-OE cells (E) compared with the Flag/HA control cells (FC >1.5; FDR <0.05). ϱ =Spearman correlation coefficient.

**Supp. Figure 6; related to Fig. 3**

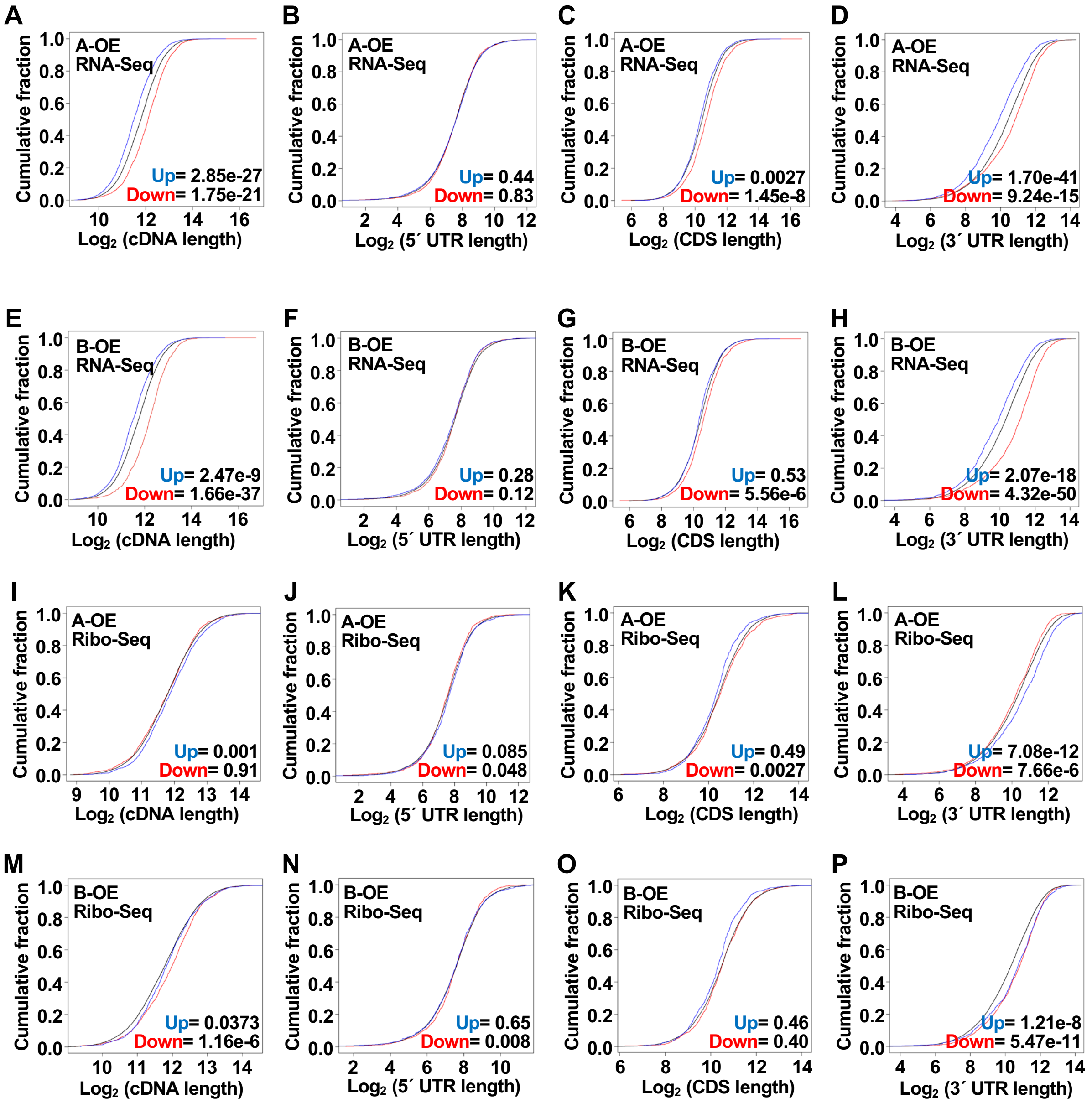


**Supplementary Figure 6. Analyses of the correlation between mRNA length and differential mRNA expression or ribosome occupancy in ZC3H7A-OE and ZC3H7B-OE HEK293 cells; Related to Figure 3.** (**A-D**) Cumulative distribution analysis of the length of the indicated regions of differentially expressed mRNAs (RNA-Seq; FC >1.5; FDR <0.05) in ZC3H7A-OE compared to control cells. (**E-H**) Cumulative distribution analysis of the length of the indicated regions of differentially expressed mRNAs in ZC3H7B-OE compared to control cells. (**I-L**) Cumulative distribution analysis of the length of the indicated regions of mRNAs with differential ribosome occupancy (Ribo-Seq; FC >1.5; FDR <0.05) in ZC3H7A-OE compared to control cells. (**M-P**) Cumulative distribution analysis of the length of the indicated regions of mRNAs with differential ribosome occupancy in ZC3H7B-OE compared to control cells. Numbers represent the p-values (Welch's Two Sample t-test) of comparisons between each group and unchanged mRNAs.

**Supp. Figure 7; related to Fig. 3**

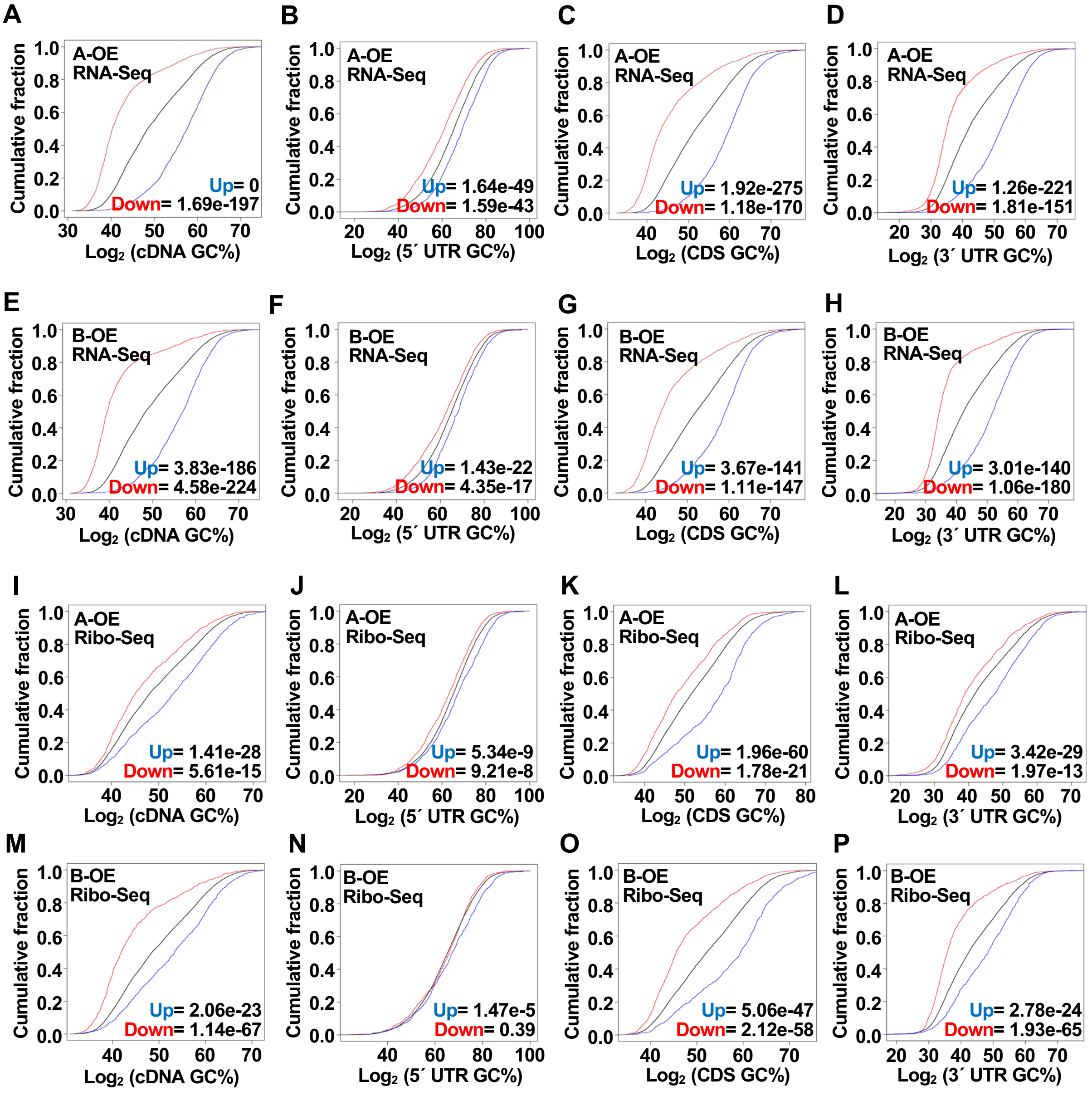


**Supplementary Figure 7. Analyses of the correlation between mRNA G/C content and differential mRNA expression or ribosome occupancy in ZC3H7A-OE and ZC3H7B-OE HEK293 cells; Related to Figure 3.** (**A-D**) Cumulative distribution analysis of percentage of G/C nucleotides in the indicated regions of differentially expressed mRNAs (FC >1.5; FDR <0.05) in ZC3H7A-OE compared to control cells. (**E-H**) Cumulative distribution analysis of percentage of G/C nucleotides in the indicated regions of differentially expressed mRNAs in ZC3H7B-OE compared to control cells. (**I-L**) Cumulative distribution analysis of percentage of G/C nucleotides in the indicated regions of mRNAs with differential ribosome occupancy (Ribo-Seq; FC >1.5; FDR <0.05) in ZC3H7A-OE compared to control cells. (**M-P**) Cumulative distribution analysis of percentage of G/C nucleotides in the indicated regions of mRNAs with differential ribosome occupancy in ZC3H7B-OE compared to control cells. Numbers represent the p-values (Welch's Two Sample t-test) of comparisons between each group and unchanged mRNAs.

**Supp. Figure 8; related to Fig. 4**

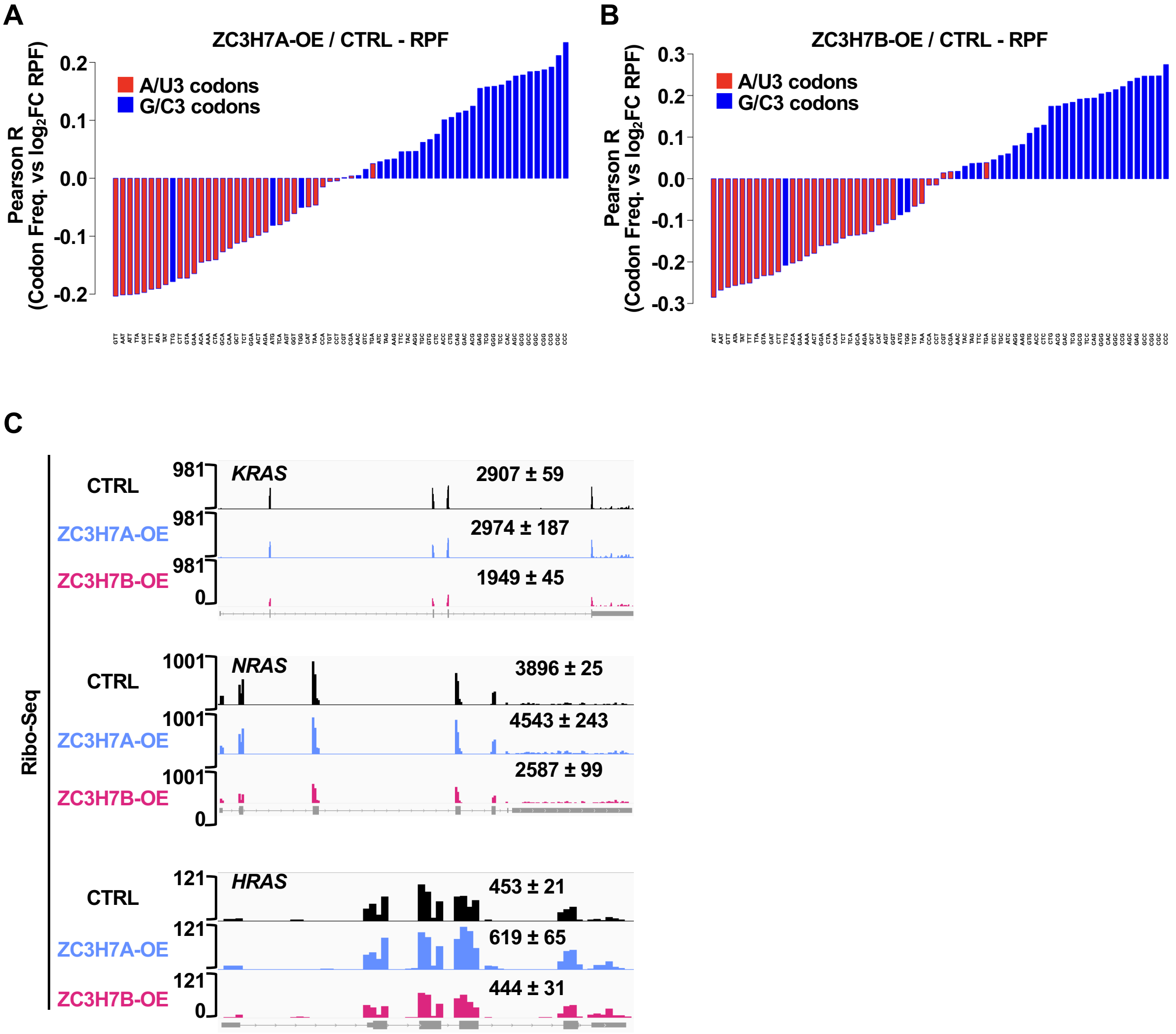


**Supplementary Figure 8. Divergent impacts of ZC3H7A/B on translation of optimal vs non-optimal mRNAs; Related to Figure 4.** (**A & B**) Waterfall plots showing Pearson correlations between codon usage frequencies and changes in ribosome occupancy in ZC3H7A-OE (A) and ZC3H7B-OE (B) cells compared with control cells. (**C**) IGV tracks visualisation of ribosome-protected fragments (RPFs) on *HRAS*, *NRAS* and *KRAS* mRNAs in the Flag/HA control, ZC3H7A-OE, and ZC3H7B-OE cells. The normalized RPF count for each gene is presented as mean ± SD.

**Supp. Figure 9; related to Fig. 5**

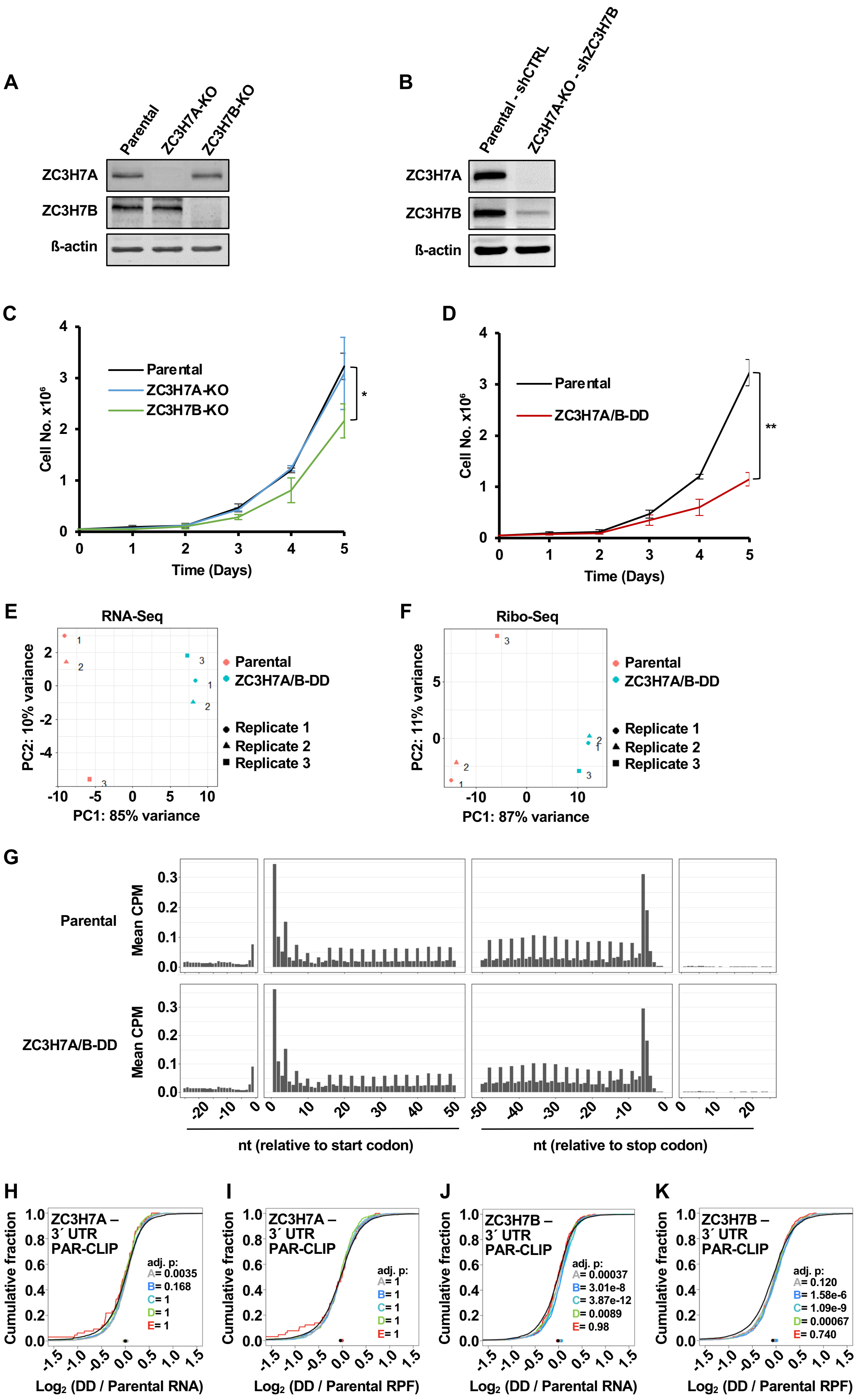

**Supplementary Figure 9. Analyses of the effects of ZC3H7A/B-double depletion on mRNA translation by Ribo-Seq; related to Figure 5. (A)** Western blot analysis with the indicated antibodies in the Parental, ZC3H7A-KO and ZC3H7B-KO HEK293 cells. **(B)** Western blot analysis with the indicated antibodies in the Parental cells expressing a non-targeting shRNA (Parental) and ZC3H7A/B–double depletion (ZC3H7A/B-DD) HEK293 cells. (**C**) Cell growth assay with Parental, ZC3H7A-KO and ZC3H7B-KO HEK293 cells at the indicated time points. Data are shown as mean ± SD; n=3 independent replicates; *p<0.05, two-tailed paired student's t-test. (**D**) Cell growth assay with ZC3H7A/B-DD and Parental cells at the indicated time points. Data are shown as mean ± SD; n=3 independent replicates; **p<0.01, two-tailed paired student's t-test. (**E & F**) PCA plot of RNA-Seq (E) and RPF (Ribo-Seq; F) of samples from Parental and ZC3H7A/B-DD cells. (**G**) Frequency of reads for footprint libraries showing the expected 3nt periodicity in relation to the translation start and stop codons in representative replicates from Parental and ZC3H7A/B-DD samples. (**H & I**) Cumulative distribution analysis of the correlation between the changes in mRNA expression (H) and ribosome occupancy (I) in ZC3H7A/B-DD compared to Parental cells and the number of binding sites across 3´ UTR for ZC3H7A. (**J & K**) Cumulative distribution analysis of the correlation between the changes in mRNA expression (J) and ribosome occupancy (K) in ZC3H7A/B-DD compared to Parental cells and the number of binding sites across 3´ UTR for ZC3H7B.

**Supp. Figure 10; related to Fig. 6**

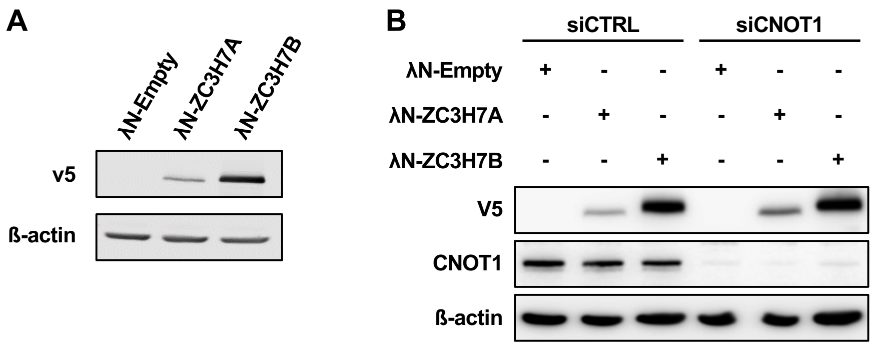


**Supplementary Figure 10. Western blot analysis of expression of λN-empty, λN-ZC3H7A, and λN-ZC3H7B; related to Figure 6.** (**A**) Western blot analysis using lysates derived from HEK293 cells transfected with *RL*-5BoxB reporter and either λN-empty, λN-ZC3H7A or λN-ZC3H7B, along with the *FL* plasmid and probed with the indicated antibodies. (**B**) Western blot analysis using lysates derived from cells transfected with siCTRL or siCNOT1 and *RL*-5BoxB reporter and either λN-empty, λN-ZC3H7A or λN-ZC3H7B, along with the *FL* plasmid and probed with indicated antibodies.

**Supp. Figure 11; related to Fig. 7**

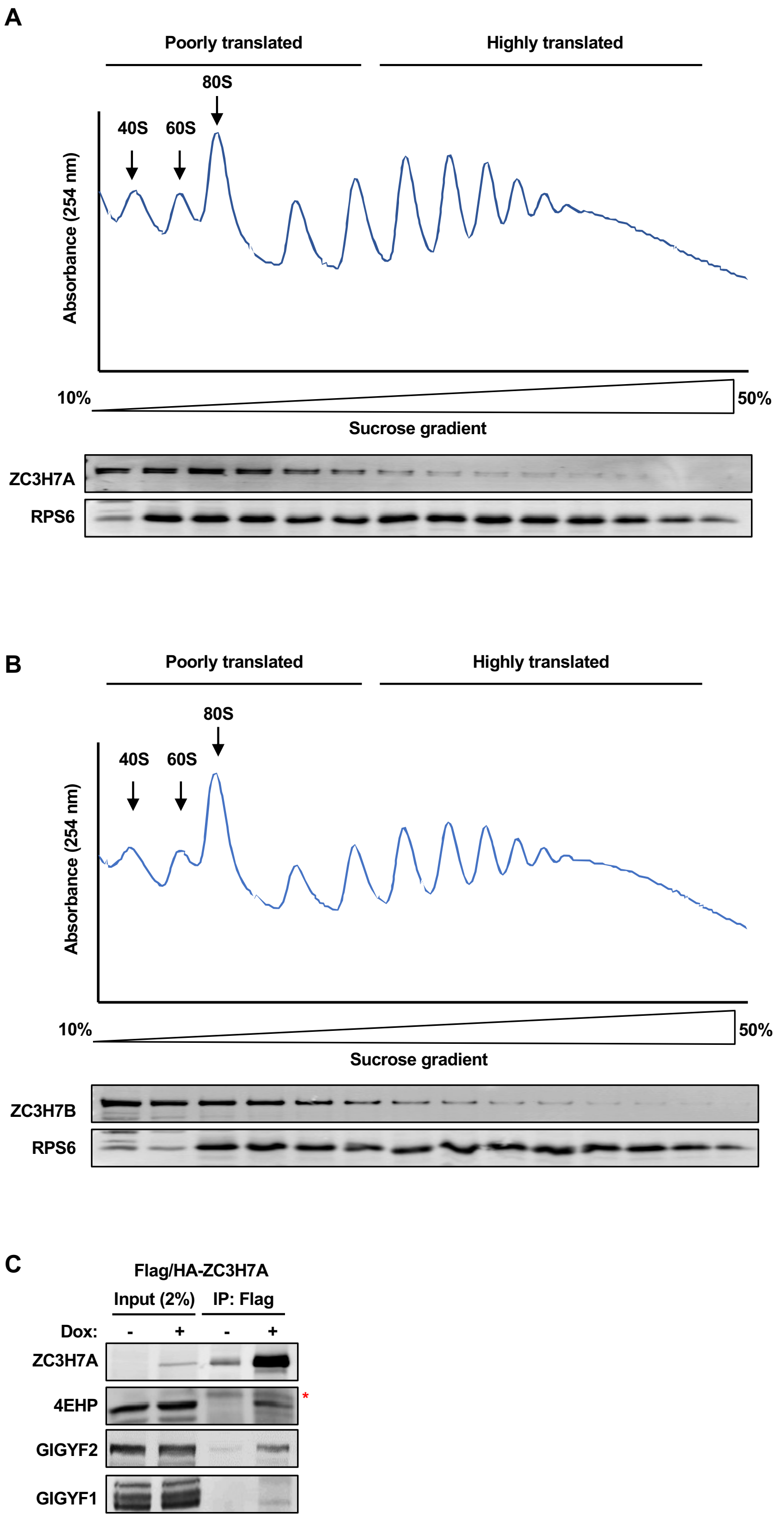


**Supplementary Figure 11. ZC3H7A/B associate with poorly translated mRNAs. related to Figure 7.** (**A**) *Top:* Polysome profiling analysis of general mRNA-ribosome association in ZC3H7A-KO HEK293 cells that express Flag/HA/ZC3H7A. *Bottom.* Western blot analysis of distribution of the ZC3H7A and ribosomal protein S6 (RPS6) in fractions derived from the sucrose gradient. (**B**) *Top:* Polysome profiling analysis of general mRNA-ribosome association in ZC3H7B-KO cells that express Flag/HA/ZC3H7B. *Bottom.* Western blot analysis of distribution of the ZC3H7B and ribosomal protein S6 (RPS6) in fractions derived from the sucrose gradient. (**C**) Co-IP for detection of interaction between Flag/HA-ZC3H7A and indicated proteins 24 h after doxycycline-induced expression of Flag/HA-ZC3H7A in ZC3H7A-KO cells.

**Supp. Figure 12**

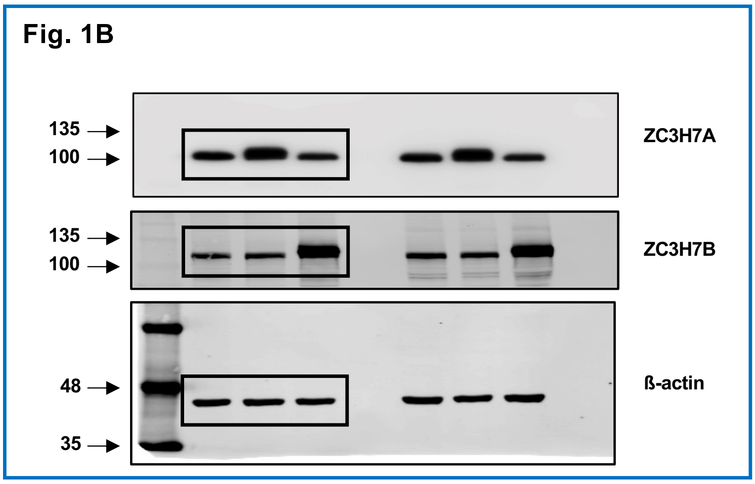


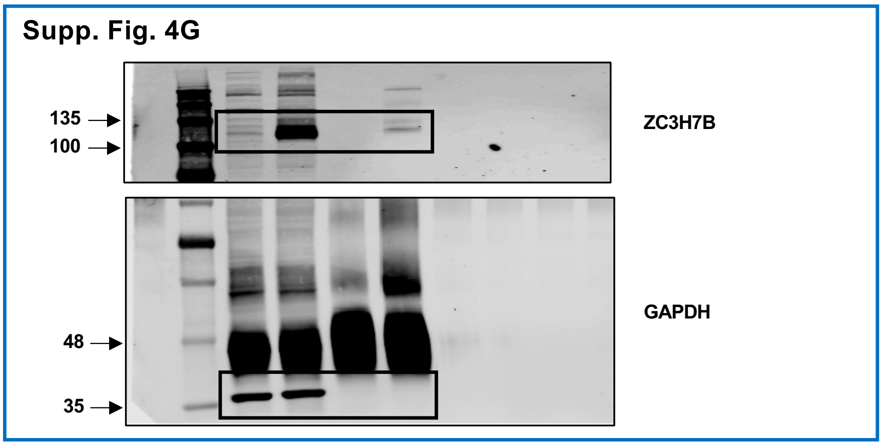


**Supplementary Figure 12.** Uncropped images of blots used in Fig. 1B and Supp. Fig. 4G.

Supp. Figure 13

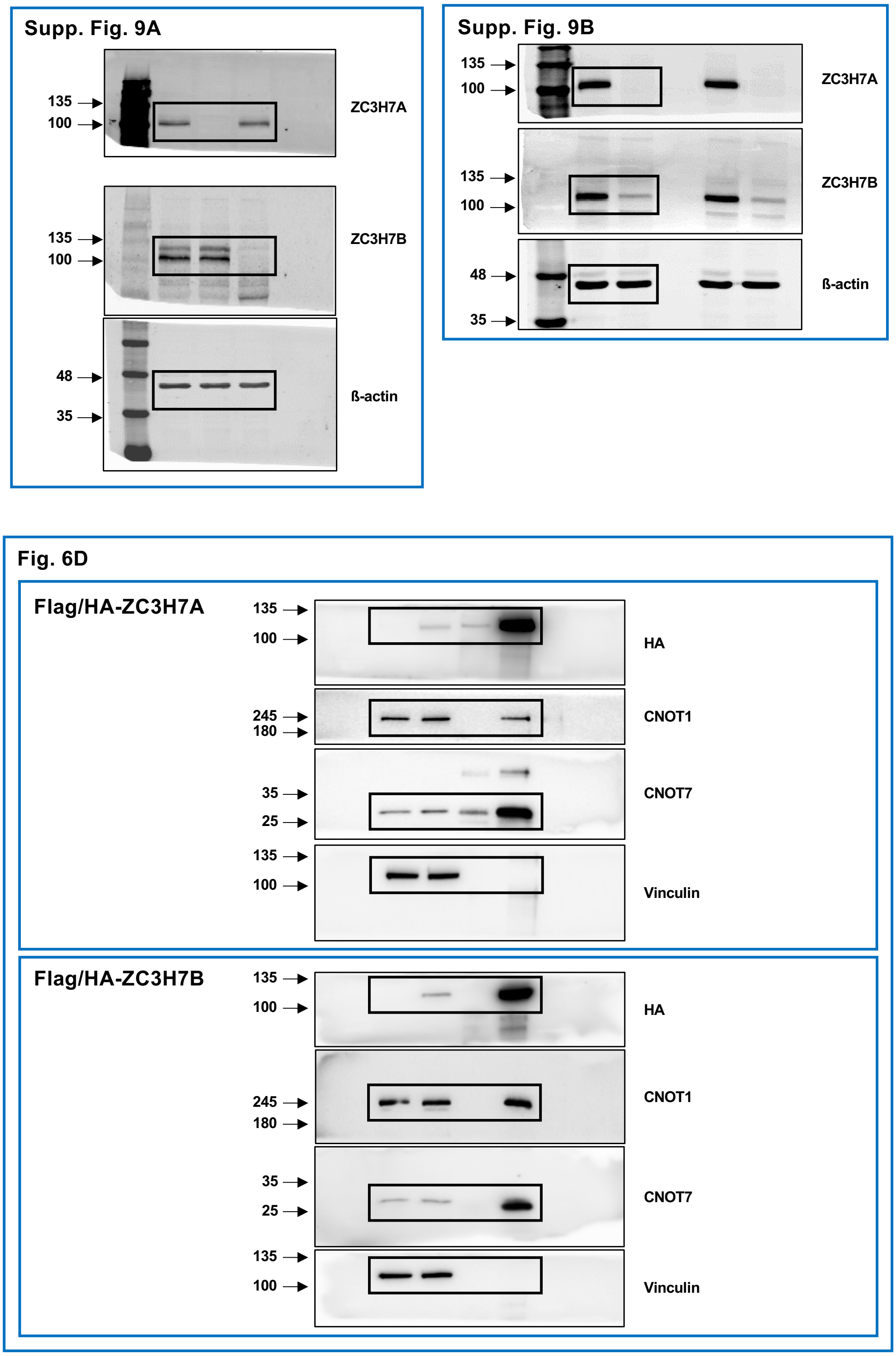


**Supplementary Figure 13.** Uncropped images of blots used in Supp. Fig. 9A & B and Fig. 6D.

### Supp. Figure 14

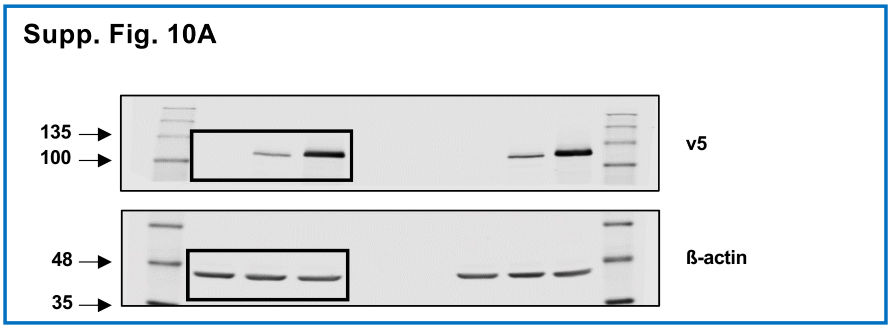


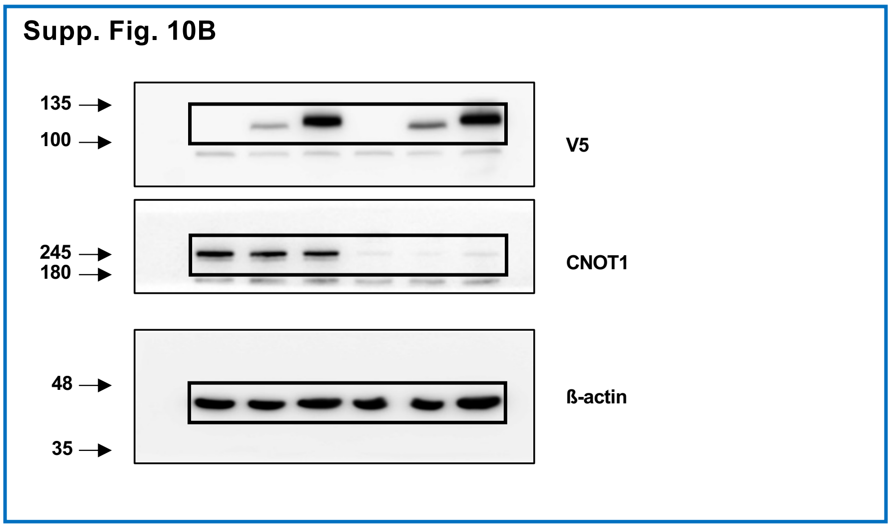


**Supplementary Figure 14.** Uncropped images of blots used in Supp. Fig. 10A & B.

Supp. Figure 15

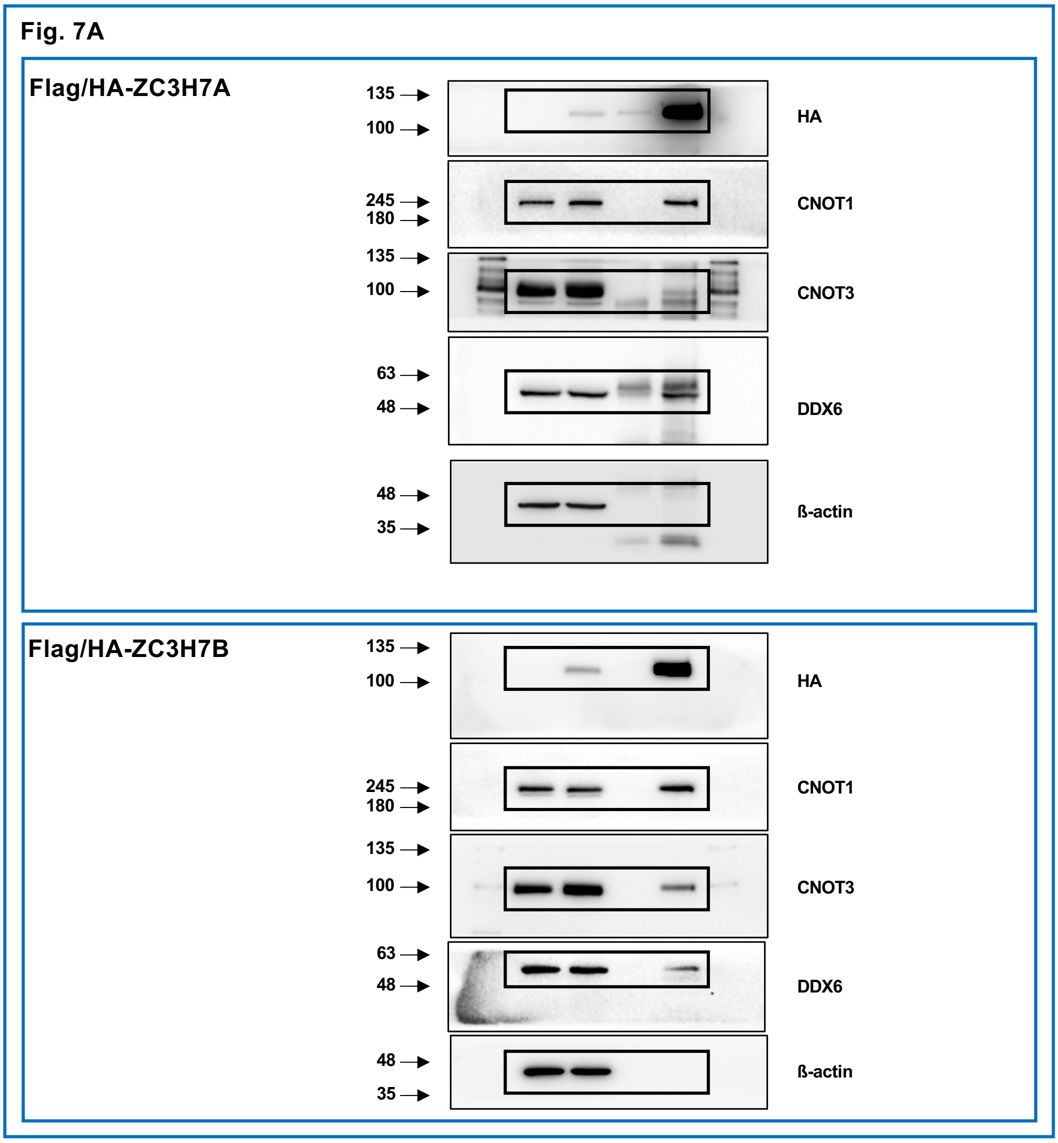


**Supplementary Figure 15.** Uncropped images of blots used in Fig. 7A.

Supp. Figure 16

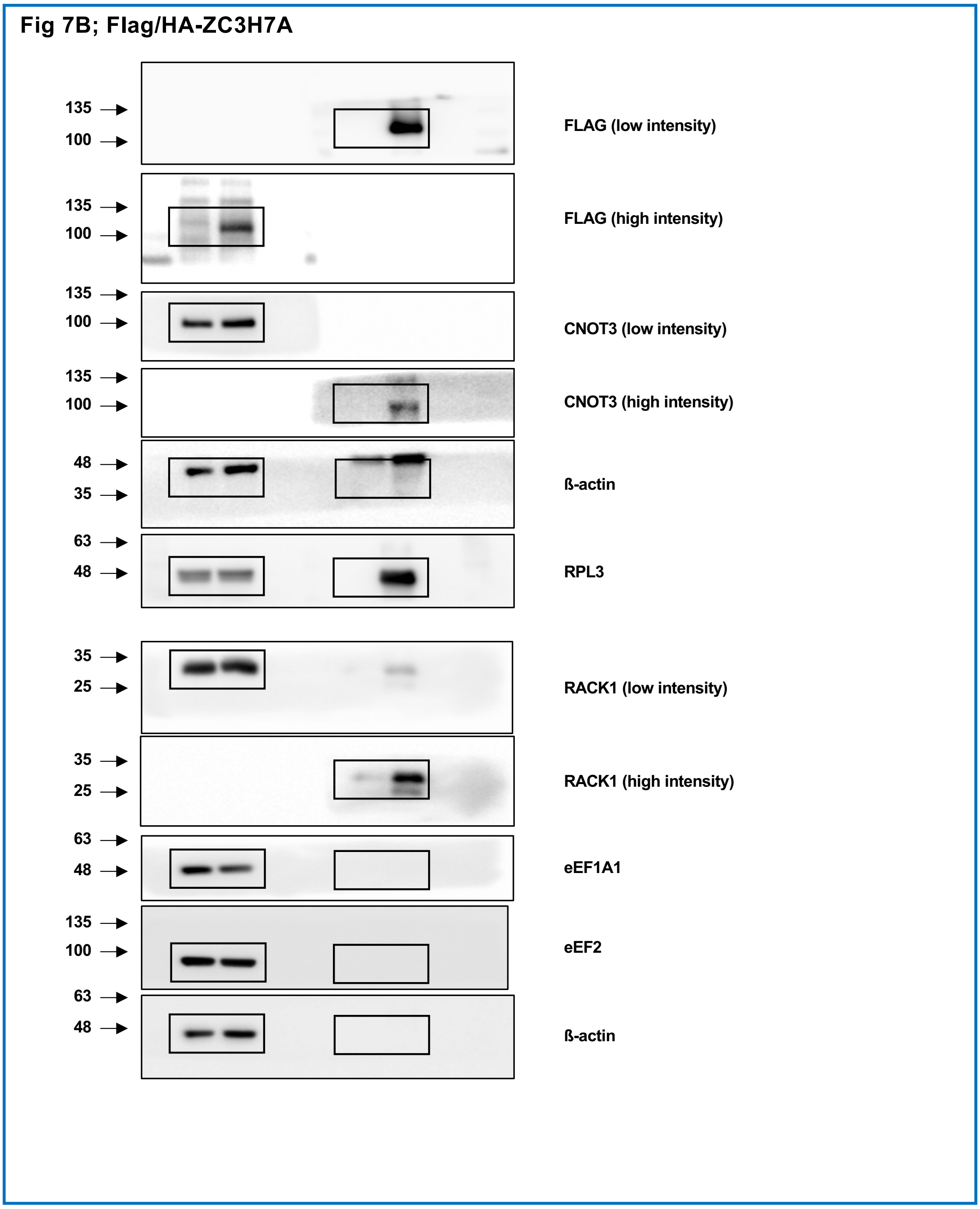


**Supplementary Figure 16.** Uncropped images of Flag/HA-ZC3H7A IP blots used in Fig. 7B.

Supp. Figure 17

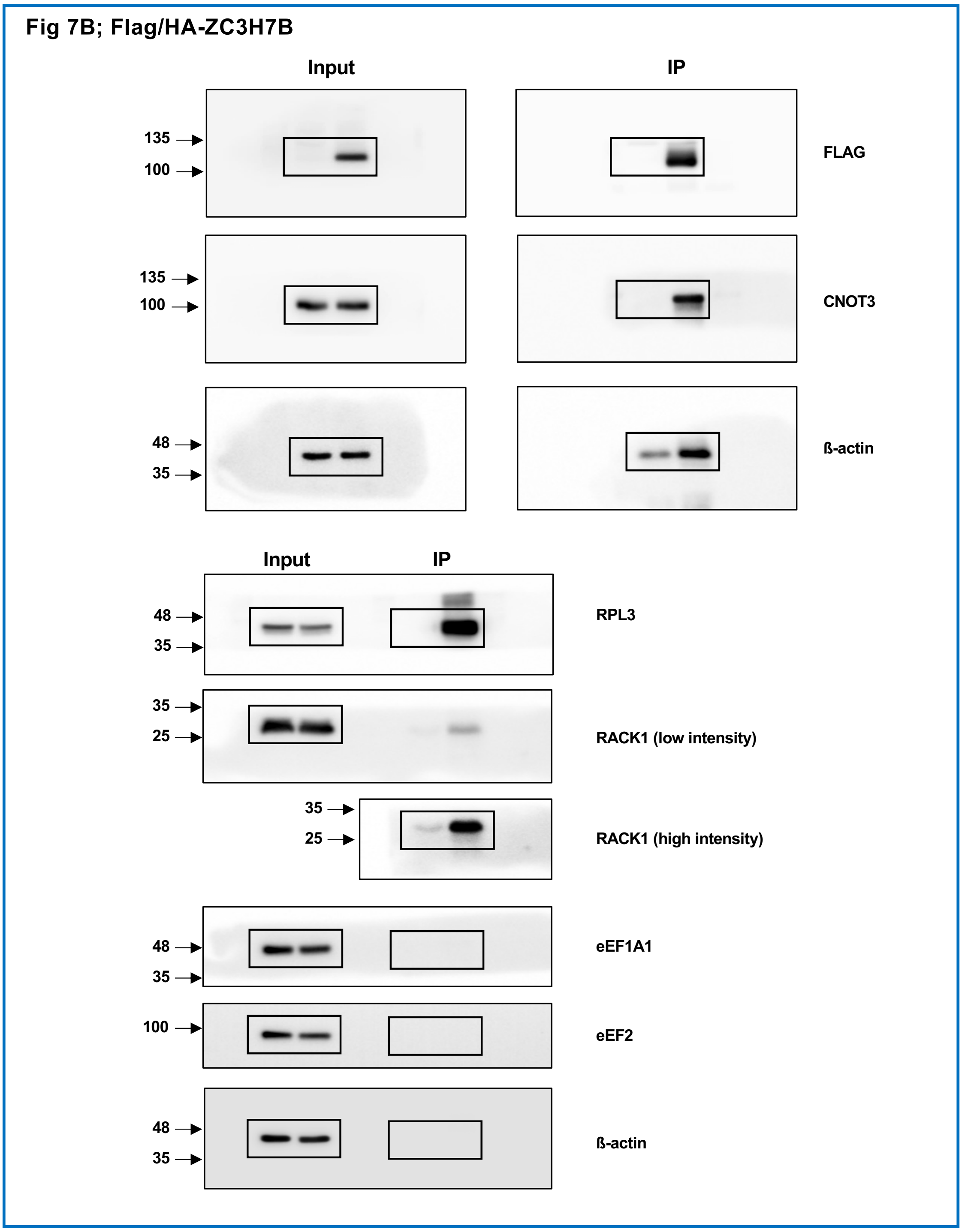


**Supplementary Figure 17.** Uncropped images of Flag/HA-ZC3H7B IP blots used in Fig. 7B.

Supp. Figure 18

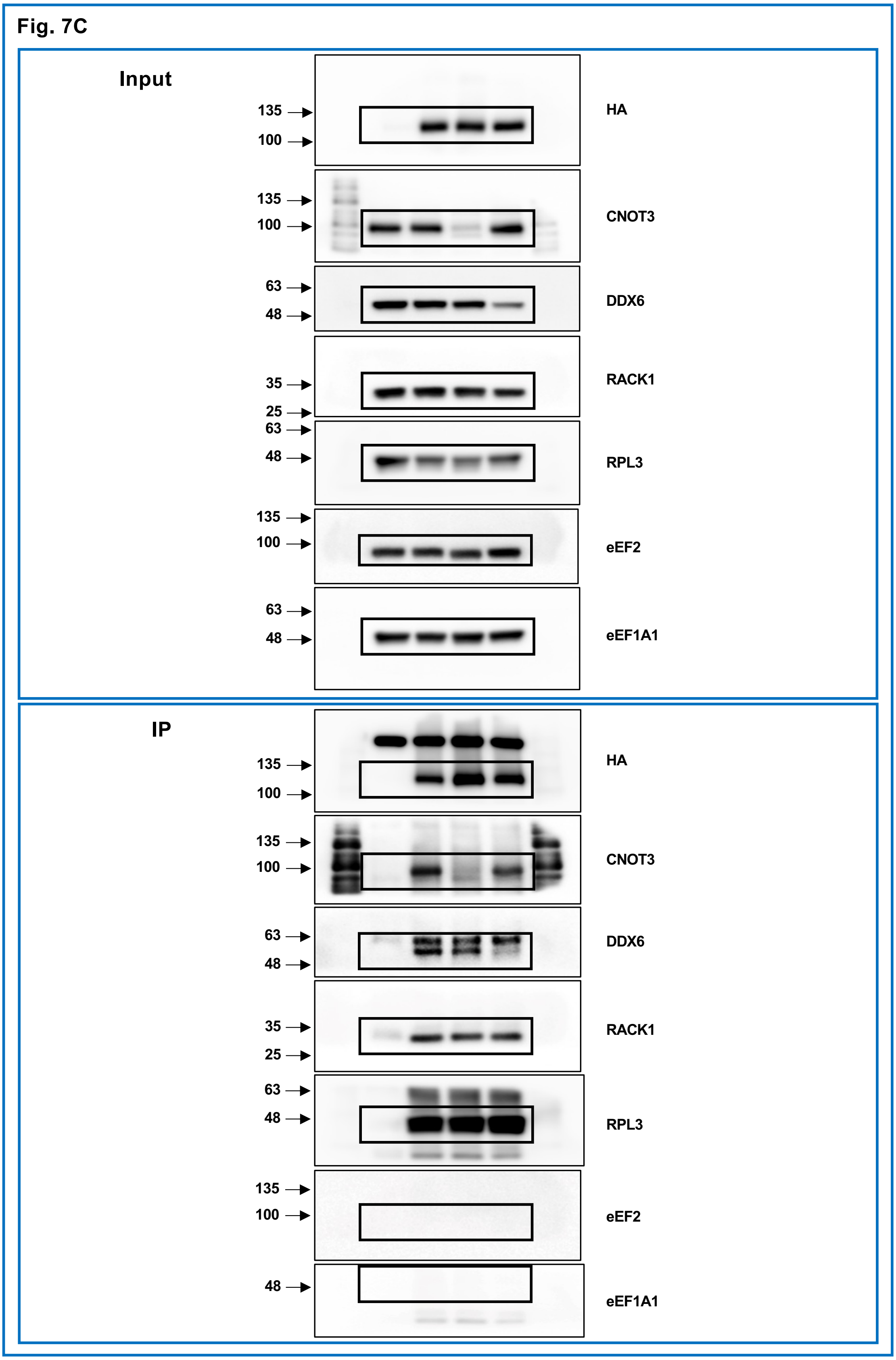


**Supplementary Figure 18.** Uncropped images of blots used in Fig. 7C.

**Supp. Figure 19**

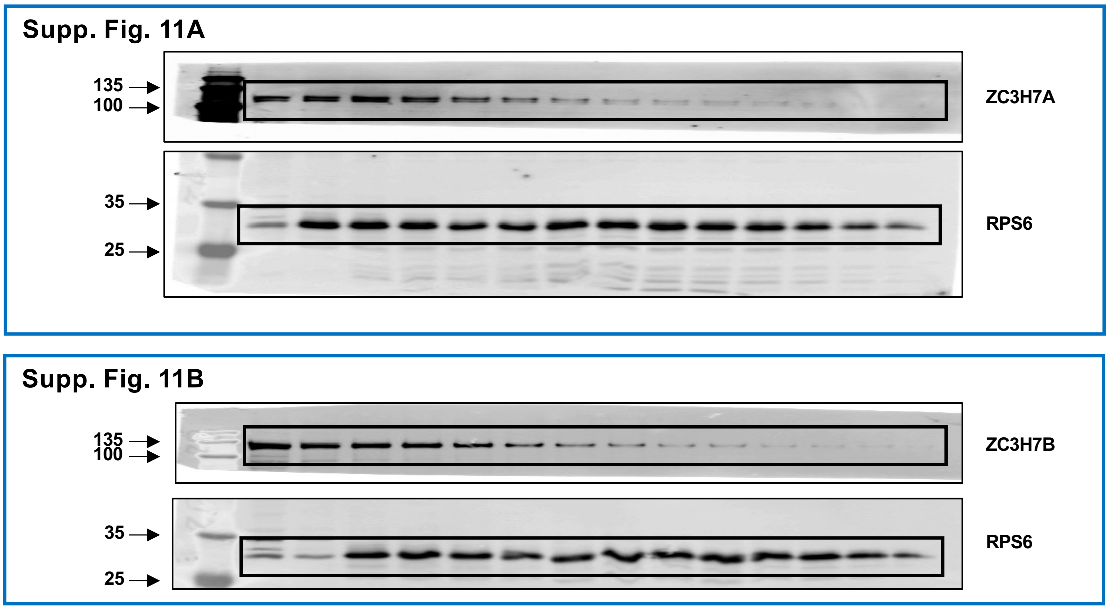


**Supplementary Figure 19.** Uncropped images of blots used in Supp. Fig. 11A & B.

**Supp. Figure 20**

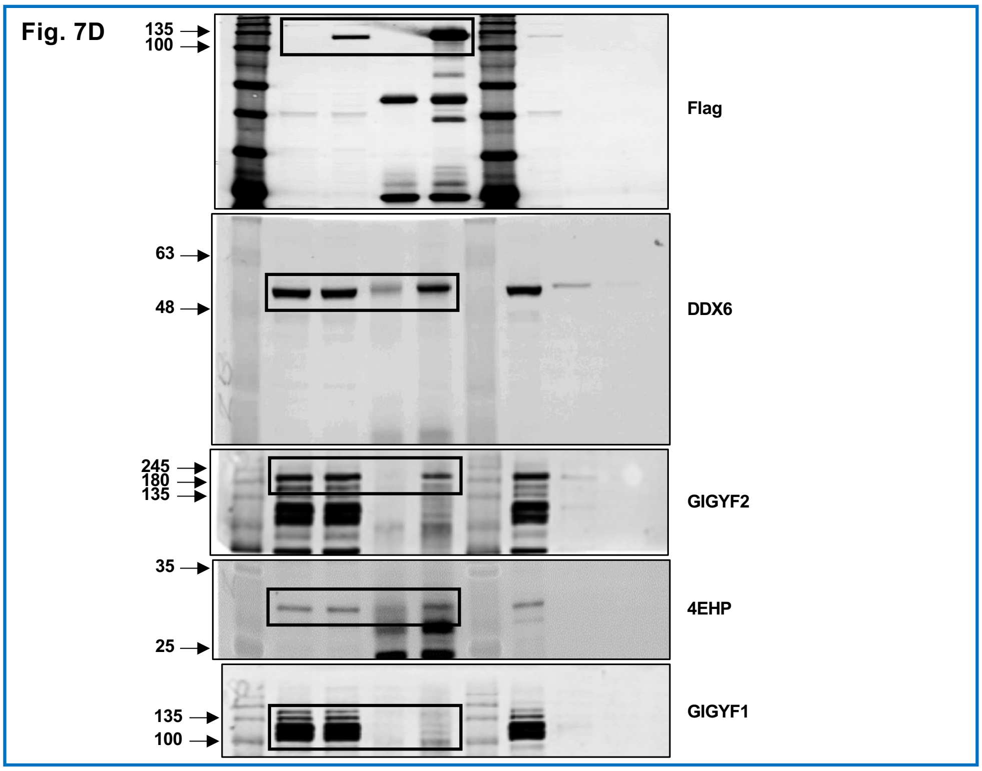


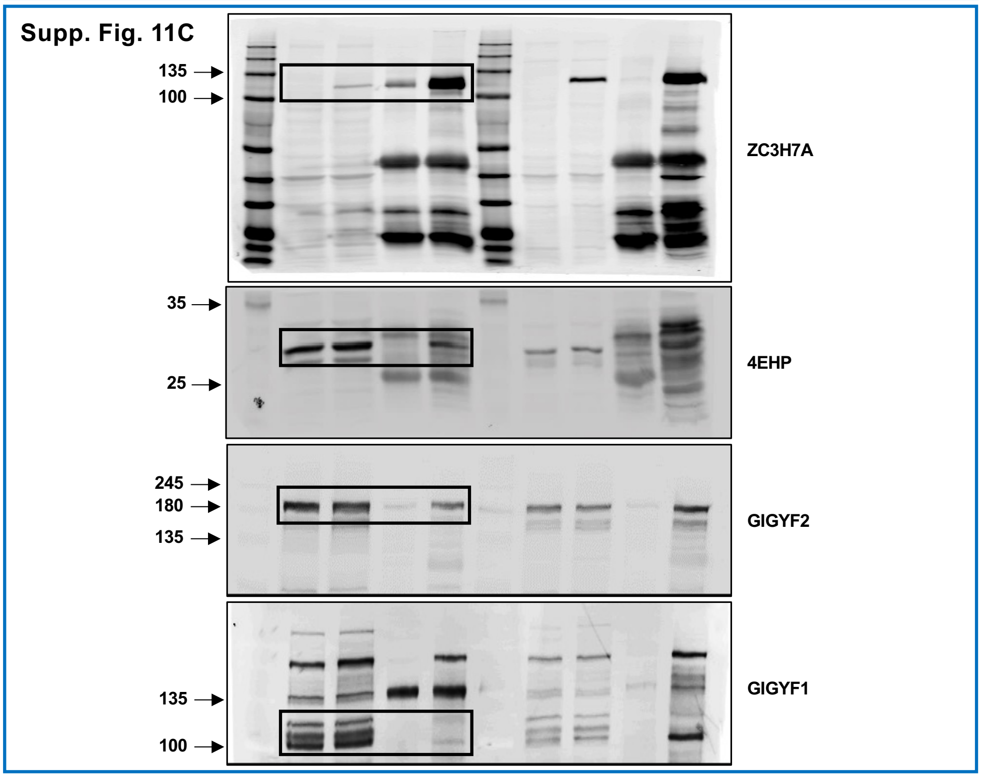


**Supplementary Figure 20.** Uncropped images of blots used in Fig. 7D & Supp. Fig. 11C.

Supp. Figure 21

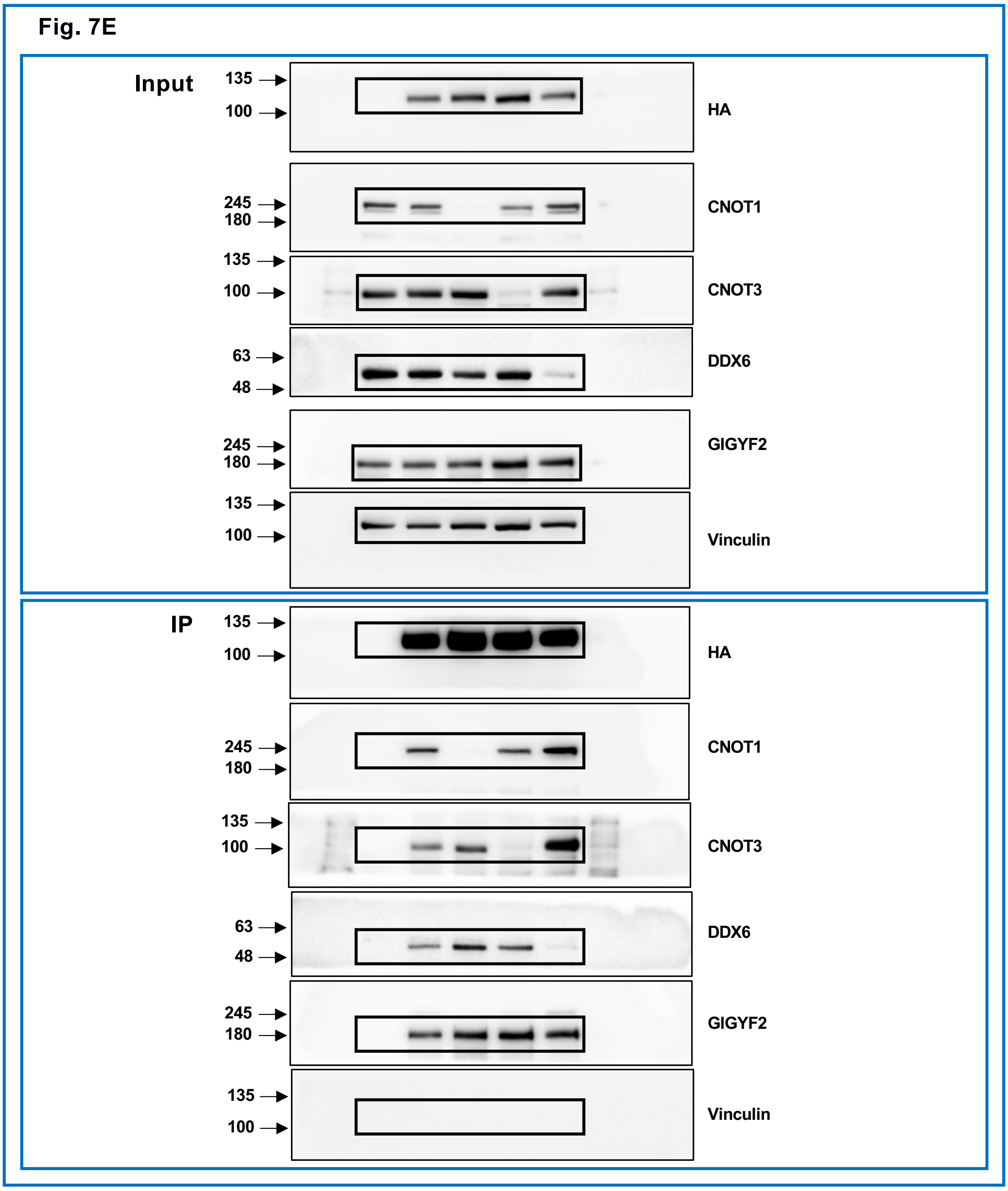


**Supplementary Figure 21.** Uncropped images of blots used in Fig. 7E.
